# Clinical severity in Fanconi anemia correlates with residual function of FANCB missense variants

**DOI:** 10.1101/772574

**Authors:** Moonjung Jung, Ramanagouda Ramanagoudr-Bhojappa, Sylvie van Twest, Rasim Ozgur Rosti, Vincent Murphy, Winnie Tan, Frank X. Donovan, Francis P. Lach, Danielle C. Kimble, Caroline S. Jiang, Roger Vaughan, Parinda Mehta, Filomena Pierri, Carlo Doufour, Arleen D. Auerbach, Andrew J. Deans, Agata Smogorzewska, Settara C. Chandrasekharappa

**Author notes:** Contributed equally to this work, in alphabetical order (Moonjung Jung, Ramanagouda Ramanagoudr-Bhojappa, Sylvie van Twest). Correspondence: Andrew J. Deans, Agata Smogorzewska or Settara C. Chandrasekharappa.

## Abstract

Fanconi anemia (FA) is the most common genetic cause of bone marrow failure, and is caused by inherited pathogenic variants in any of 22 genes. Of these, only *FANCB* is X-linked. We describe a cohort of 19 children with *FANCB* variants, from 16 families of the International Fanconi Anemia Registry (IFAR). Those with *FANCB* deletion or truncation demonstrate earlier than average onset of bone marrow failure, and more severe congenital abnormalities compared to a large series of FA individuals in the published reports. This reflects the indispensable role of FANCB protein in the enzymatic activation of FANCD2 monoubiquitination, an essential step in the repair of DNA interstrand crosslinks. For *FANCB* missense variants, more variable severity is associated with the extent of residual FANCD2 monoubiquitination activity. We used transcript analysis, genetic complementation, and biochemical reconstitution of FANCD2 monoubiquitination to determine the pathogenicity of each variant. Aberrant splicing and transcript destabilization was associated with two missence variants. Individuals carrying missense variants with drastically reduced FANCD2 monoubiquitination in biochemical and/or cell-based assays showed earlier onset of hematologic disease and shorter survival. Conversely, variants with near-normal FANCD2 monoubiquitination were associated with more favorable outcome. Our study reveals a genotype-phenotype correlation within the FA-B complementation group of FA, where severity is linked to the extent of residual FANCD2 monoubiquitination.

**KEY POINTS:**

- X-linked *FANCB* pathogenic variants predominantly cause acute, early onset bone marrow failure and severe congenital abnormalities
- Biochemical and cell-based assays with patient variants reveal functional properties of FANCB that associate with clinical severity

## INTRODUCTION

Fanconi anemia (FA) is an inherited disorder caused by DNA repair deficiency, which results in progressive bone marrow failure leading to aplastic anemia. FA is also associated with congenital abnormalities and a highly elevated risk of acute myeloid leukemia and head and neck cancers.^1^ In a large series of patients, the average age at onset of hematological abnormalities was reported to be 7.6 years,^2^ and about two-thirds of patients exhibit one or more congenital abnormalities.^3^ At least 22 FA-causing genes have been identified, named *FANCA-FANCW*,^4^ but only *FANCB* (MIM# 300515) is X-linked recessive.^5^ This makes it especially important to identify *FANCB* variants as to pathogenicity, and whether *de novo* or not, in order to provide proper genetic counseling. Variants in *FANCB* account for ~4% of all male cases of FA,^6^ and are often associated with severe congenital abnormalities resembling VACTERL (<u>V</u>ertebral abnormalities, <u>A</u>nal atresia, <u>C</u>ardiac defects, <u>T</u>racheoesophageal fistula, <u>E</u>sophageal atresia, <u>R</u>enal abnormalities or <u>L</u>imb abnormalities) association with hydrocephalus.^7,8^ The course of hematological disease and severity of VACTERL-H or any genotype-phenotype correlations among *FANCB* individuals remains to be elucidated.

FANCB protein is a component of the FA core complex that also contains the products of six other FANC genes (A, C, E, F, G and L) together with <u>FA</u>-<u>a</u>ssociated <u>p</u>roteins (FAAP100 and FAAP20).^9^ The FA core complex is necessary to signal the existence of stalled replication forks, caused by DNA interstrand crosslinks (ICL), covalent bonds between Watson and Crick DNA strands. It is required for monoubiquitination of the DNA binding proteins FANCD2 and FANCI, which form the ID2 heterodimer. In particular, FANCB, FANCL, and FAAP100 are the critical minimal components required for *in vitro* monoubiquitination of ID2.^10,11^ While FANCL is the RING E3 ligase necessary for ubiquitination to occur, FANCB and FAAP100 play an enigmatic role in the reaction.

Very few studies have explored the function of the FANCB protein, despite its identification as a critical enzymatic component of the FA core complex. This is in part because the protein contains no known predicted domains, and also because it is intractable to work with in isolation. It was shown that co-expression and purification of a recombinant complex of FANC<u>B</u>, FANC<u>L</u>, and FAAP<u>100</u> (BL100 complex) can overcome these difficulties,^10–12^ and this reflects a similar interdependence of stability for these three proteins in human cells.^5,13,14^ Recombinant BL100 complex is twice its predicted mass, reflective of a dimerization of the complex through the FANCB subunit.^11^ BL100 can also interact independently with the CEF (FANC<u>C</u>, FANC<u>E</u>, FANC<u>F</u>) and AG20 (FANC<u>A</u>, FANC<u>G</u>, FAAP<u>20</u>) subcomplexes of the FA core complex. Here, through biochemical and cell-based assays we demonstrate FANCB’s central role in the FA core complex assembly and activity, and establish how patient-associated variants disrupt DNA damage signaling in FA. We present molecular diagnosis, functional evaluation of the variants and clinical presentations of the largest cohort of individuals with *FANCB* variants yet described. We find that individuals with missense variants, in general, had a delayed onset of hematologic disease, and longer overall survival, which correlate with the extent of residual function retained by the variant FANCB protein.

## METHODS

### Study participants

The study subjects include individuals diagnosed with FA and their family members enrolled in the International Fanconi Anemia Registry (IFAR), following written informed consent/assent. The Institutional Review Board of the Rockefeller University, New York, USA, has approved these studies. The Office of Human Subjects Research at the National Institutes of Health and Institutional Review Board of the National Human Genome Research Institute (NHGRI) approved the reception of de-identified cell lines and DNA samples from The Rockefeller University and analysis of the underlying molecular variants.

### Identification of disease-causing FANCB variants

Genomic DNA was isolated from peripheral blood, fibroblast, or EBV-immortalized lymphoblastoid cell line (LCL). Targeted Nextgen sequencing, Sanger sequencing, array comparative genomic hybridization (aCGH) and SNP arrays were employed to identify sequence variants as described earlier,^15^ and further outlined in supplemental methods.

### Reverse transcription-PCR (RT-PCR)

Total RNA from individual-derived cell lines was obtained using RNeasy Plus Mini Kit (Qiagen). Complementary DNA (cDNA) synthesis was done using SuperScript™ III or IV First-Strand Synthesis System (Thermo Fisher Scientific). Screening for aberrant *FANCB* splice products was done as described further in supplemental methods.

### Quantitative fluorescence PCR (qf-PCR)

A modified version of the previously described qf-PCR method was used to measure the relative levels of aberrant splice or missense product vs normal splice product.^16,17^ A FAM-labeled primer (M13F-FAM) was included to generate fluorescently labeled products. Primers to generate splice products with similar sizes were designed and are listed in Table S1. Details of PCR reactions are provided in supplemental methods.

### Cell culture

Fibroblasts were cultured in DMEM (Gibco) plus 15% FBS (Atlanta Biologicals), 1% Pen Strep (Gibco), 1% GlutaMAX™ (Gibco) and 1% MEM non-essential amino acids (Gibco). LCLs were cultured in RPMI 1640 (Gibco) plus 20% FBS, 1% Pen Strep and 1% GlutaMAX™. Fibroblast cell lines were transformed and/or immortalized by expression of HPV16 E6E7 and the catalytic subunit of human telomerase (hTERT), respectively.

### Complementation of FANCB deficient cell lines with wild type (WT) or variant FANCB

The WT *FANCB* and missense variants cloning, lentiviral vector preparation, and complementation experiments were performed as described earlier,^18^ and further outlined in supplemental methods.

### Recombinant FANCB protein co-purification

*FANCB* variant was cloned into pFL-FANCB-FANCL-FAAP100, and purified protein was generated from resultant baculovirus as described in Swuec et al^12^ and summarized in the supplemental methods.

### In vitro ubiquitination experiments

All protein purifications and reactions were performed exactly as described in van Twest et al^11^ and summarized in supplemental methods, substituting variant FANCB-containing BL100 complexes where appropriate. StrepII-FANCD2 and Flag-FANCI were detected with αStrepII tag (ab76949, Abcam) or αFlag-tag (OAEA00002, Aviva Systems Biology) antibodies.

### Statistical analyses of clinical data

The statistical analyses of clinical data for time to death and time to hematologic disease were performed as outlined in supplemental methods.

## RESULTS

### A wide spectrum of inherited or de novo FANCB pathogenic variants cause FA

In a comprehensive effort towards molecular diagnosis of FA, we identified disease-causing *FANCB* variants in 19 individuals from 16 families enrolled in the IFAR (Figure 1A, Table 1). The IFAR families were of varied race/ethnicity, including Caucasian, Hispanic and African-American. Table 1 describes the clinical information and genotype of these 19 individuals from the IFAR and two additional individuals (# 9 and 21) published previously by others.^8,19^ Among the 19 IFAR individuals, we observed two large deletions, 590kb (individuals 1-2) and 520kb (individual 3) encompassing the *FANCB* gene in two families. SNP arrays and aCGH showed that these deletions included the entire *FANCB* gene as well as an adjacent upstream gene, *MOSPD2*, and a downstream gene, *GLRA2* (Figure S1). Further, we identified sequence variants predicted to generate a truncated protein in 9 individuals, including two variants affecting splice junctions (individuals 4-5), three resulting in a nonsense codon (individuals 6-8), and four indels from three families (individuals 11-14). Individual 15 carried two overlapping *de novo* variants, a missense (NM_001018113:c.2249G>T; p.G750V) and an indel (NM_001018113:c.2249_2252delGAAG p.G750Vfs*5). The proportion of missense variant was observed to go down from 80% in blood to 53% in the LCL, suggesting positive growth selection for cells with indel variant (Table S2). Five individuals from four families carried missense variants (individuals 16-20). All missense variants appeared in highly conserved residues across 99 FANCB orthologs (Figure 1B, Table S3). Additionally, we previously reported individual #10,^18^ who carried an unstable 9154 bp intragenic duplication, exhibiting mosaicism in both the proband and the carrier mother; this individual is not discussed further in this report. It is notable that each of the families in this study had a unique pathogenic variant, with no evidence for a common founder.

**Figure 1.**
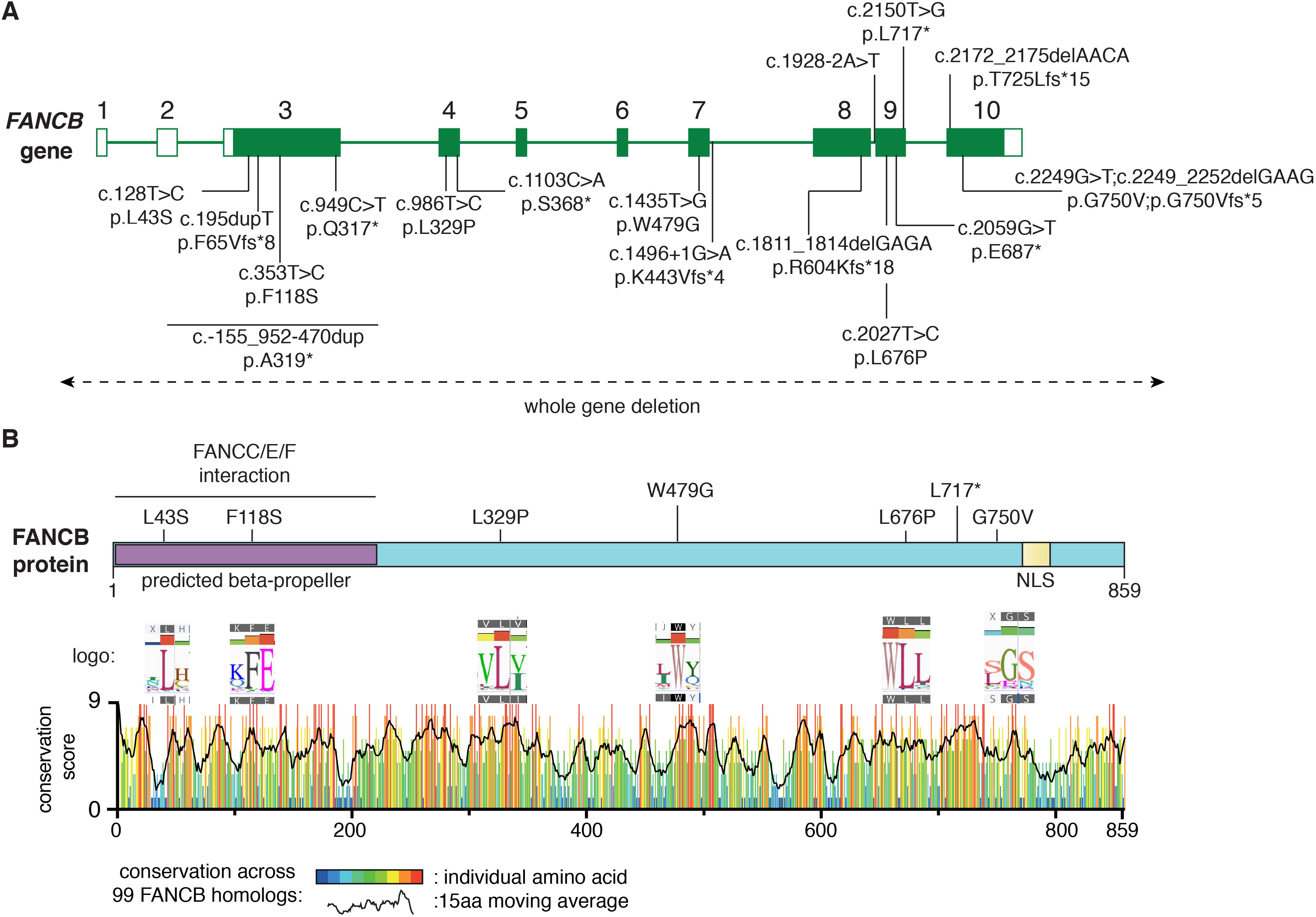
Schematic of *FANCB* gene and protein with identified variants. (A) The *FANCB* gene with intron-exon organization, along with the cDNA coordinate and the predicted amino acid changes for each variant from 16 IFAR families we identified are indicated. One IFAR family with a duplication c.-155_952-470dup (p.A319*) has been reported earlier^18^ and the duplicated region is shown as a solid line. Two variants from published reports, F118S^19^ and L717*,^8^ are indicated here as well, as their functional status are being evaluated in this study. The whole gene deletion (WGD) extending beyond *FANCB* are indicated by a dotted line with arrows at the end. For the splice junction variant in intron 7, c.1496+1G>A, the predicted protein change is based on RNA analysis, while lack of RNA prevented identifying the consequences on the protein of the intron 8 variant, c.1928-2A>T. (B) Schematic of FANCB protein indicating location of missense variants. The predicted structural motifs, beta propeller and NLS are shown. The conservation score plot of FANCB across 99 vertebrate homologs is shown below. For each amino acid that is altered in an FA patient, the “sequence logo” derived from 99 FANCB vertebrate homologs is shown for the affected amino acid and the adjacent ones. The consensus sequence and the human WT FANCB sequence are above and below the logo, respectively. The location of the nonsense variant p.L717* is also shown.

Availability of parental DNA for 13/16 IFAR families allowed us to infer the origin of the pathogenic variant. Nine were inherited (12 individuals) while four were presumed *de novo* (i.e. arose during gametogenesis in a parent, or very early in embryogenesis) (Table 1). We performed deep sequencing of the variant region in the proband and parents using MiSeq method (Table S4). At a read depth in the range of 223,000-609,000, we confirmed the variants to be absent in the parents and *de novo* in the probands (Table S2). Despite being *de novo* in origin, blood mosaicism was not observed and the variant frequency was ~100% in three individuals. In the fourth individual (#15) there were two overlapping *de novo* variants. Finally, through deep sequencing, we confirmed that mosaicism was not present in any of the five probands with inherited variants or their parents/siblings (Table S5).

### FANCB RNA analysis reveals aberrant splice and unstable transcripts

Availability of cell lines for all individuals except #5 (c.1928-2A>T) allowed us to analyze the splicing and stability of *FANCB* transcripts. Aberrant splicing products were observed in individuals 4, 7 and 19. Individual 7, carrying a nonsense variant (c.949C>T; p.Q317*) near the end of exon 3, generated a second transcript that eliminates 74 bp from the end of exon 3 (p.G294Cfs*3) (Figure 2A). Individual 4, carrying a canonical splice variant (c.1496+1G>A), generated a frameshift transcript due to skipping of exon 7 (170 bp; p.K443Vfs*4) (Figure 2B). Individual 19, carrying a missense variant in exon7 (c.1435T>G; p.W479G), generated an additional transcript without exon 7 (Figure 2B). The relative levels of transcripts with and without exon 7 were 13.5% and 86.5%, respectively (Figure 2C-E).

**Figure 2.**
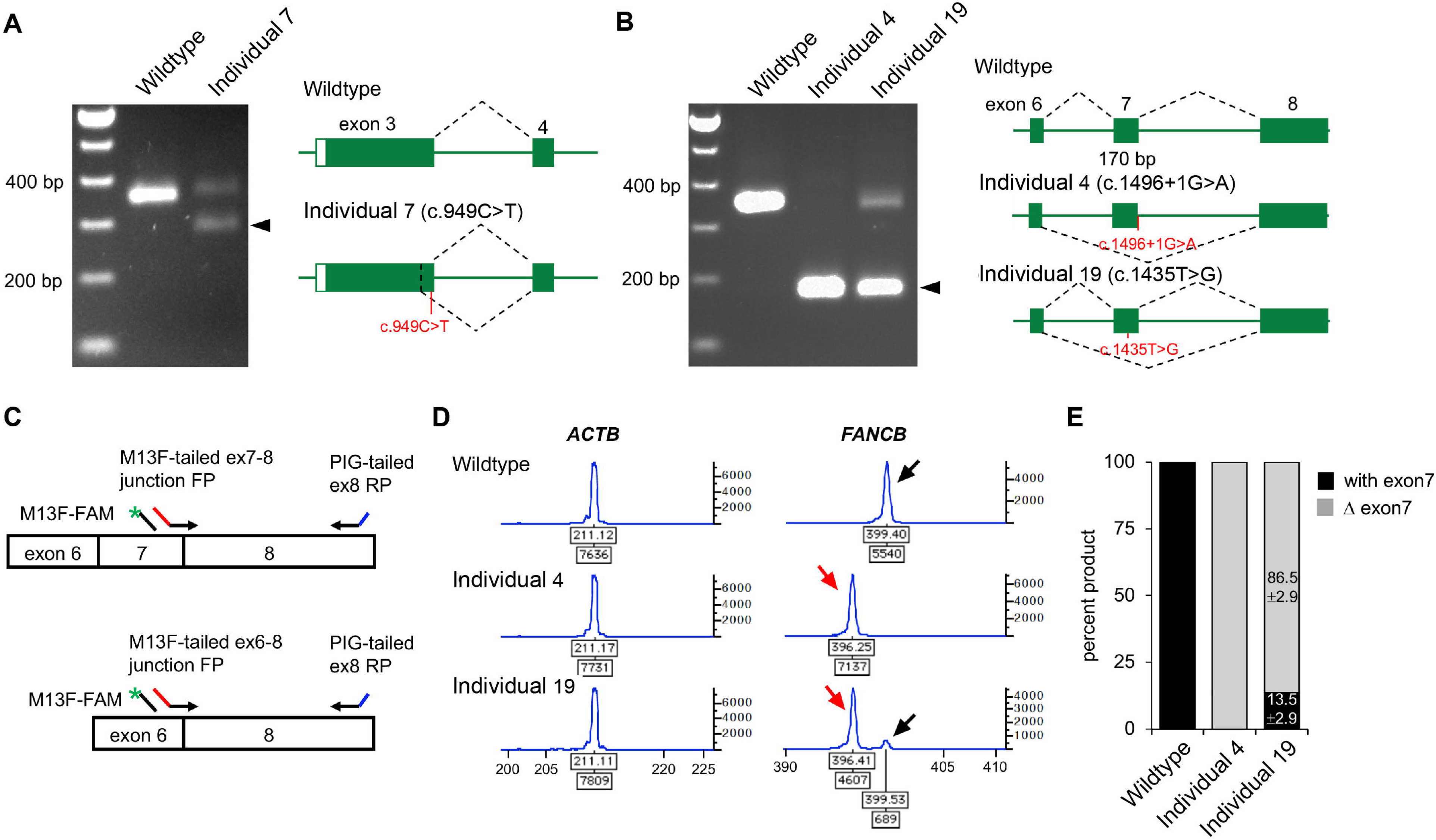
Aberrant *FANCB* splice products and their quantitation. (A) RT-PCR analysis of fibroblast cell line RNA from individual 7 with c.949C>T (p.Q317*) variant in exon 3. The gel shows, along with the predicted size product, an additional smaller product (arrow) that lacked the last 74 bp from the end of exon 3 (c.878_951del74; p.G294Cfs3*). Both products encode truncated proteins. RNA from a fibroblast cell line from an unaffected individual is used as WT control. (B) RT-PCR analysis of fibroblast cell lines RNA from individuals 4 and 19 carrying splice junction c.1496+1G>A and a missense c.1435T>G (p.W479G) variants, respectively. The individual 4 RNA shows exclusively exon 7 skipping (c.1327_1496del170; p.K443Vfs*4) (arrow). Individual 19 also results in exon 7 skipping along with a normal size product. (C) Schematics of qf-RT-PCR for relative quantification of transcripts with and without exon 7. Two *FANCB* specific forward primers that bind to either exon 7-8 junction or exon 6-8 junction and a reverse primer that bind to exon 8 were designed to amplify products from transcripts with (top) or without exon 7 (bottom), respectively. The product with exon 7 is 3 bp longer than that from without exon 7. FAM-labeled M13 forward primer (M13F-FAM) was included to generate fluorescently labeled products. FP; forward primer. RP; reverse primer. (D) Representative qf-RT-PCR profiles from individuals 4 and 19. Product size and intensity are on the x- and y-axis respectively; both numbers are shown under each peak. The left panel shows products from *ACTB*, an internal transcript control. The right panel shows *FANCB* products. The fragment size for products with exon 7 (black arrow) and without exon 7 (red arrow) appeared around 399 bp and 396 bp, respectively. Data from WT, individual 4 and individual 19 are presented on the top, middle and bottom rows respectively. (E) Percentage of transcripts with or without exon 7, average from eight independent assays.

The quantitative analysis of the LCL from individual 15 revealed the presence of indel variant transcripts only, while gDNA analysis showed the presence of both the indel and missense variants (Figure S2A-B), suggesting that the missense variant transcript is highly unstable. The unstable nature of this variant may be the reason behind the observed positive growth selection for cells with the indel variant from blood to LCL (Table S2).

### Functional evaluation of constructs expressing missense variants in a *FANCB* null cell line

Unlike the non-functional nature of a truncated protein variant, the effect of a missense variant is difficult to determine without functional evaluation. We assessed the function of all the identified *FANCB* missense variants using both patient-derived cell line and by overexpression of the variant in an FANCB null cell line. Cell lines from all individual with missense variants lacked FANCD2 ubiquitination upon MMC treatment, confirming the absence of a functional FA core complex (Figure S3A-B). Cell line from individual 15, who carried both a missense and an indel variant, showed low level of FANCD2 ubiquitination. However, FANCD2 ubiquitination was completely restored upon complementation with WT FANCB expression (Figure S3A) identifying the FANCB variants as causative for FA phenotypes in these individuals.

We next introduced cDNA expression constructs into an FANCB null cell line, and evaluated FANCB function using three assays post-treatment with MMC: FANCD2 ubiquitination, FANCD2 foci formation, and cell viability (Figure 3). FANCD2 ubiquitination was nearly absent in the cell line expressing the L43S variant, minimal in the L329P and G750V cell lines, and comparable to WT in the W479G and L676P cell lines (Figure 3A). The FANCD2 foci formation assay (Figure 3B-C) and cell survival assay (Figure 3D) also revealed variable activity of the variants: L43S showed very little function, L329P was moderate, and W479G and L676P displayed substantial activity. The G750V-complemented cell line showed little function in these assays, however we found that the lentiviral transcript carrying c.2249G>T (G750V) variant was expressed at 3.5-fold lower level than WT FANCB transcript (Figure S2C). This resulted in reduced expression of G750V protein, indicating that the c.2249G>T variant destabilizes the transcript. Except for G750V, which shows globally low expression of HA-FANCB, the remaining missense variants also had reduced nuclear localization (Figure S4).

**Figure 3.**
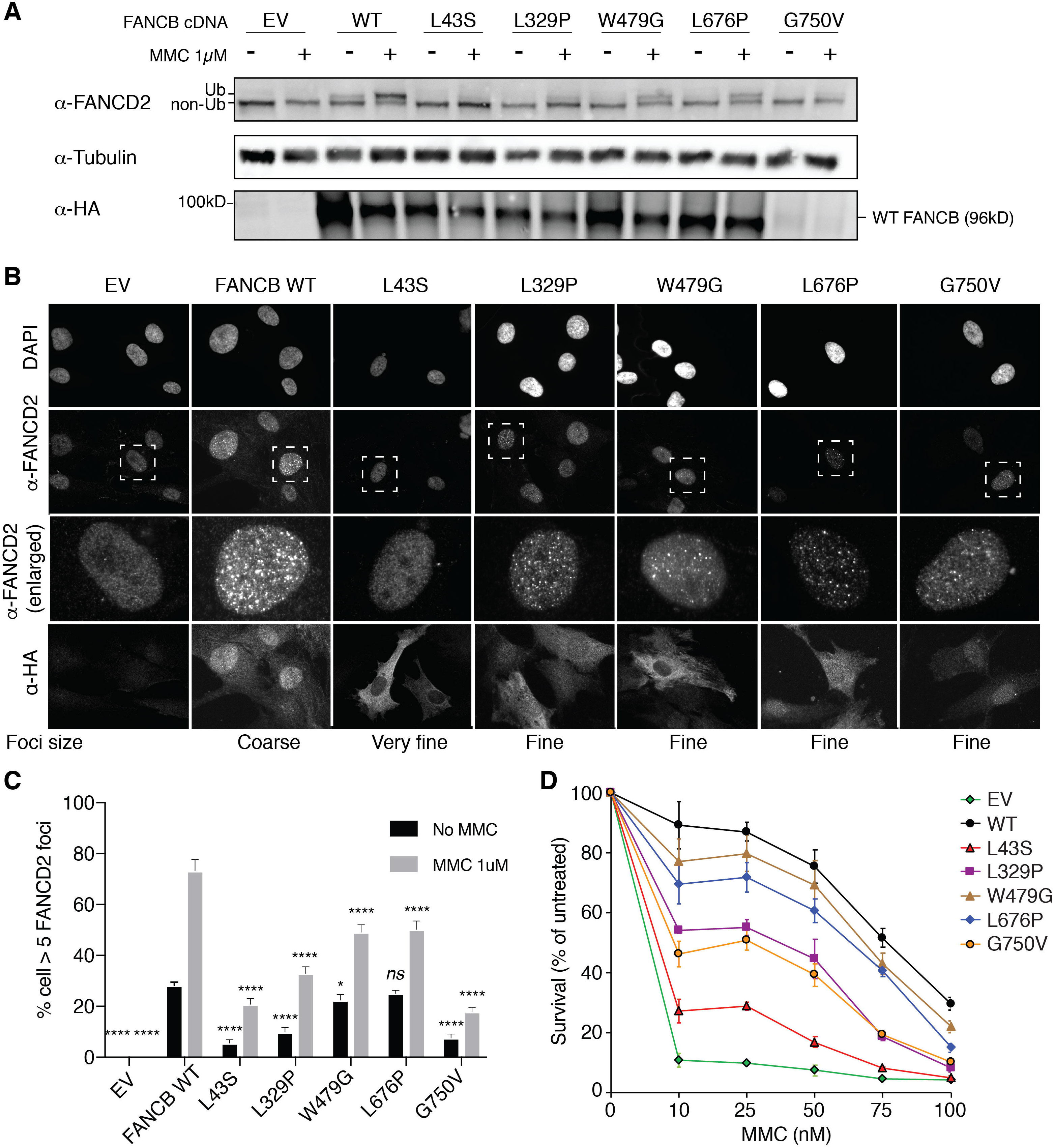
Overexpression of missense variants in FANCB-null fibroblasts show varying degree of residual function. HA-tagged cDNA of *FANCB* WT or five missense variants were overexpressed by lentiviral vector in FANCB-null fibroblasts, followed by functional analysis. (A) Western blot of FANCD2, tubulin and HA-FANCB. Near-normal FANCD2 ubiquitination was observed upon MMC exposure with expression of p.W479G and p.L676P, while expression of p.L43S showed almost no FANCD2 ubiquitination. Expression of p.L329P and p.G750V resulted in low level of FANCD2 ubiquitination. Of note, HA-FANCB p.G750V was unstable. Tubulin was used as loading control. (B) Immunofluorescence of FANCD2 and HA-FANCB. Smaller-sized and decreased number of FANCD2 foci in cells expressing HA-FANCB missense variants were apparent compared with HA-FANCB WT. (C) Quantification of FANCD2 foci-positive cells. One hundred cells were counted in triplicate per each experiment. Three independent experiments were performed. Statistical analysis was performed using one-way ANOVA followed by Dunnett’s multiple comparison test. * *P* < 0.05 and **** *P* < 0.001. *ns*: not significant (D) Fibroblasts expressing FANCB WT or missense variants were treated with corresponding doses of MMC after which surviving cells were counted and survival was calculated relative to untreated condition. Three independent experiments were performed. A graph from a representative experiment is shown.

### Efficiency of missense variants in recombinant FA core complex assembly and in vitro *FANCD2:FANCI* ubiquitination

To define how FANCB variants directly affected the assembly of the FA core complex and subcomplexes, we utilized an *in vitro* reconstitution assay and biochemical test of FA core complex assembly and activity in ubiquitination of FANCD2 and FANCI proteins. In addition to the missense variants described above, we included F118S, the only other missense variant reported in the literature,^19^ and L717*, the truncation variant missing only the last 142 amino acids.^8^ The FA core complex is made up of three subcomplexes: BL100, AG20 and CEF.^20^ Baculoviral expression constructs of FA core complex proteins were used for co-expression of either BL100 subcomplex proteins or all the nine proteins of the FA core complex (Figure 4A). Flag-tagged WT and missense variant FANCB protein were used as bait to purify protein complexes, resolved using PAGE, and then the components were quantitated. The assembly of the BL100 subcomplex was not affected by the missense variants, nor the 142 amino acid C-terminal truncation variant (L717*) (Figure 4B). However, the L43S and F118S variants showed significantly reduced AG20 and CEF association (Figure 4C). Using FFAS-3D,^21^ this region of the protein was predicted to have structural homology to other proteins containing beta-propellar structures, a known protein:protein interaction motif. SMURFLite analysis, which predicts beta-propellar structures more stringently,^22^ concurred with high confidence (Figure S5). The remaining variants, which lie outside of this region, L329P, G750V, and L676P and the L717* truncation, retained WT levels of FA core complex assembly (Figure 4C).

**Figure 4.**
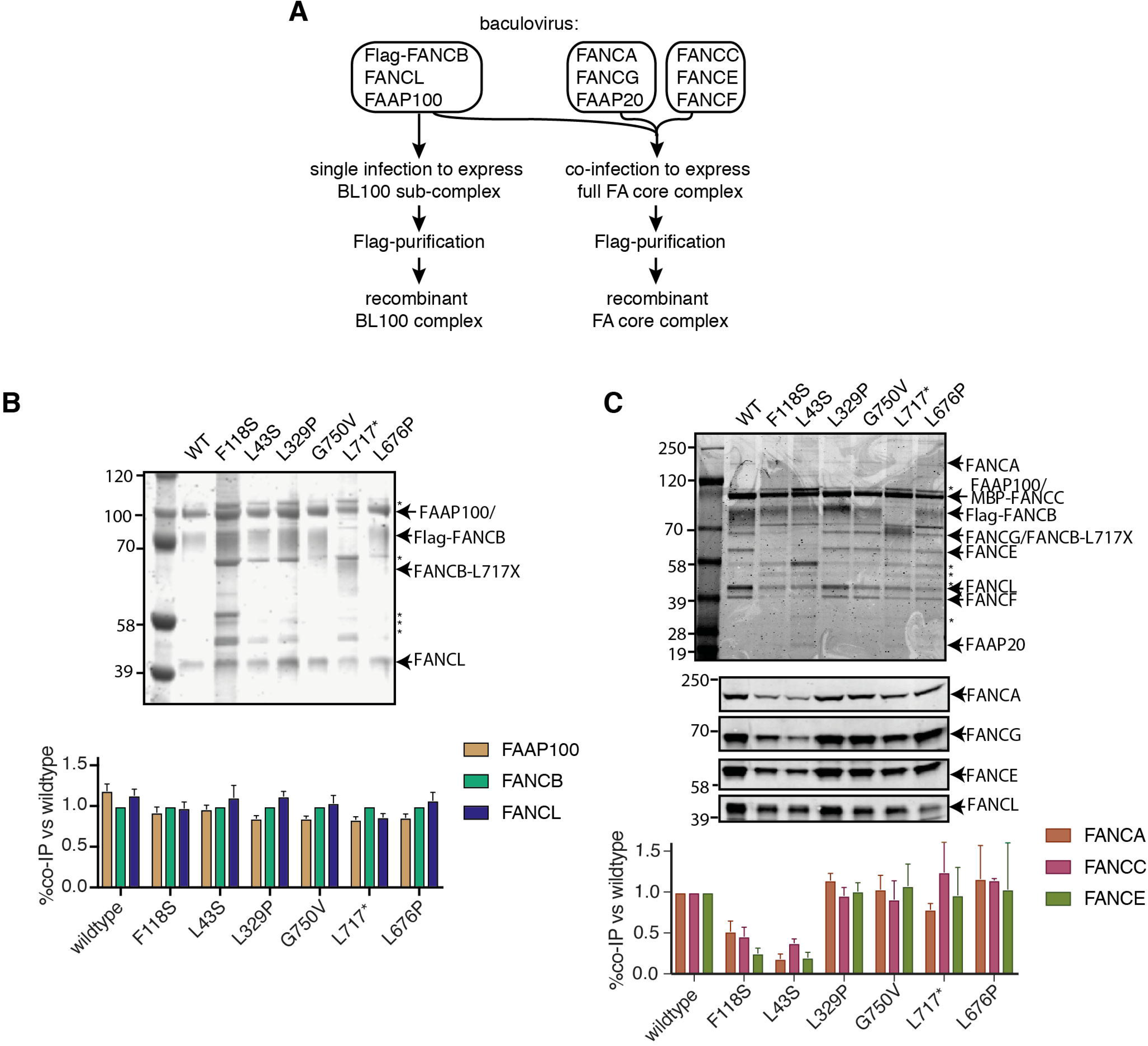
N-terminal FANCB variants affect FA core complex assembly. (A) Schematic for protein purifications of human FANCB complexes purified from baculovirus infected insect cells. (B) Coomassie blue stained SDS-PAGE gel of affinity purified FANCB-FANCL-FAAP100 (BL100) complex and bar graph (n=2 experiments) indicating relative molar amount of FANCL and FAAP100 co-purified with WT of variant Flag-FANCB protein as bait. (C) As in B but using full FA core complex. Western blots for FANCA, FANCG, FANCE, and FANCL are shown to identify individual Coomassie-stained bands. Asterisks represent contaminant proteins that bind to Flag-affinity resin.

Because several variant BL100 complexes failed to properly interact with the remainder of the core complex, we evaluated their effect on *in vitro* ubiquitination of FANCD2 and FANCI using BL100 complexes purified in isolation and then added to other core complex components (Figure 5A). The levels of monoubiquitylation of FANCD2 and FANCI were measured using western blot analysis. As predicted by poor FA core complex assembly, the L43S and F118S variants showed reduced FANCD2 and FANCI monoubiquitination (Figure 5B). As previously observed, the kinetics of FANCI monoubiquitination was slower than that for FANCD2, and the N-terminal changes effected both reactions proportionally. The *in vitro* ubiquitination activity of complexes with L329P and L676P variants resembled WT FANCB for FANCD2 monoubiquitination, however, these had a slower rate of FANCI monoubiquitination (Figure 5C). Unexpectedly, given that L717* still forms an intact FA core complex, this truncation variant was completely defective in both FANCD2 and FANCI monoubiquitination, even after a 90 minute incubation period in which equivalent WT and missense variants had attained 100% FANCD2 monoubiquitination. The G750V variant showed near-normal levels of monoubiquitination of both FANCD2 and FANCI, indicating that this variant at protein level may not contribute towards pathogenicity.

**Figure 5:**
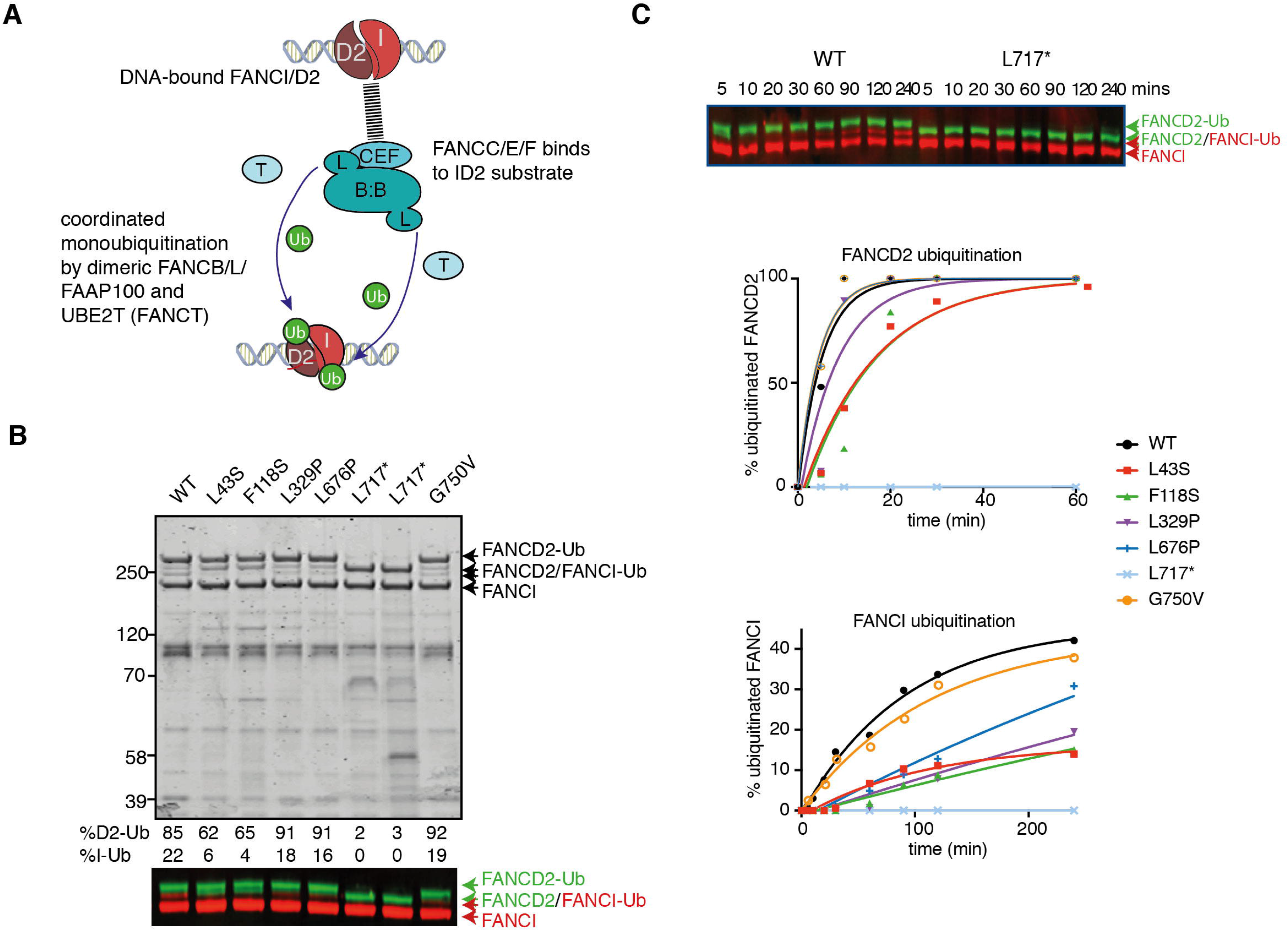
Effect of FANCB patient variants on ubiquitination activity of the FA core complex. (A) Schematic of *in vitro* ubiquitination reaction (B) Results of an example monoubiquitination assay: Coomassie stained SDS-PAGE gel to compare WT BL100 to BL100 with FANCB patient variants in 90min ubiquitination reaction. Western blotting of FANCD2 and FANCI reveals the extent of ubiquitinated protein was calculated from western blots using anti-StrepII-Fancd2 (green) or anti-Flag-FancI (red) (FANCI). Two different preparations of L717* complex are shown. (C) Example of time course experiment comparing monoubiquitination activity of WT and L717*-FANCB containing reactions by western blot (above) or quantified ubiquitinated forms of FANCD2 and FANCI. Similar results were obtained from n=4 experiments.

### Individuals with FANCB pathogenic variants exhibit multiple congenital anomalies, early hematological disease onset and earlier death

Congenital malformations resembling VACTERL-H association were seen in 15/19 (79%) individuals with *FANCB* variants with the predominant features being renal (R) and limb (L) abnormalities (observed in 13 individuals each) (Table 1), which is consistent with previous observations.^7,23^ In general, developmental defects were multiple and severe for the individuals in our study: termination of pregnancy elected for three IFAR individuals diagnosed prenatally, and for one previously reported in the literature, and survival was limited to 0.033, 0.2 and 0.5 month for three other individuals. Interestingly, the three individuals in our cohort without VACTERL-H malformations did not carry truncation variants, which suggests that the residual function of the FA core complex may protect against the most severe developmental defects (Table 1).

The median survival time and mean age of onset of hematological disease have been reported as 24 years^24^ and 7.6 years,^2^ respectively. Our data suggest that FA-B individuals have earlier death with a median time to death of 104 months (n=15 including individual 21; Table 1, Figure S6A), and earlier onset of hematological disease with a median and mean age of onset at 26 months and 33.6 months, respectively (n=11 including individual 21; Table 1, Figure S6B).

Seven individuals underwent hematopoietic stem cell transplant (HSCT) for their hematologic disease; two died of HSCT complications, while the other five are still alive at 108, 132, 132, 168, and 168 months of age. The survival for the remaining four with no HSCT was 1.1, 80, 104, and 136 months. In all, excluding the seven individuals who underwent HSCT, the survival data, which was available for 8 individuals, revealed that no one survived beyond 136 months, and the median survival was 15.05 months.

To assess the genotype and phenotype correlation, we grouped variants based on the severity of structural alterations in the FANCB protein (WGD/truncation vs. missense). The median time to death was earlier in the WGD/truncation variants group (15.05 months; range 0.03 to 168 months) than in the missense group (104 months; range 45 to 132 months), but was not statistically significant (Log-Rank test = 0.5198, p-value = 0.4709; HR = 1.693 (95% CI: 0.398 - 7.198)) (Figure 6A). Similarly, the median time to the hematologic disease was earlier in the WGD/truncation variant group (6.25 months; range 0.25 to 48 months), than in the missense group (43 months; range 12 to 102 months), but was also not statistically significant (Log-Rank test = 2.3962, p value = 0.1216; HR = 3.058 (95% CI: 0.669 - 13.973)) (Figure 6B). To assess the effect of residual function of missense variants on clinical outcome, we plotted outcomes of the activity of the identified variants and the clinical outcome in the individuals with these variants (Figure 6C). Although the number of individuals is small, clinical outcomes appear better with increased residual function of FANCB.

**Figure 6.**
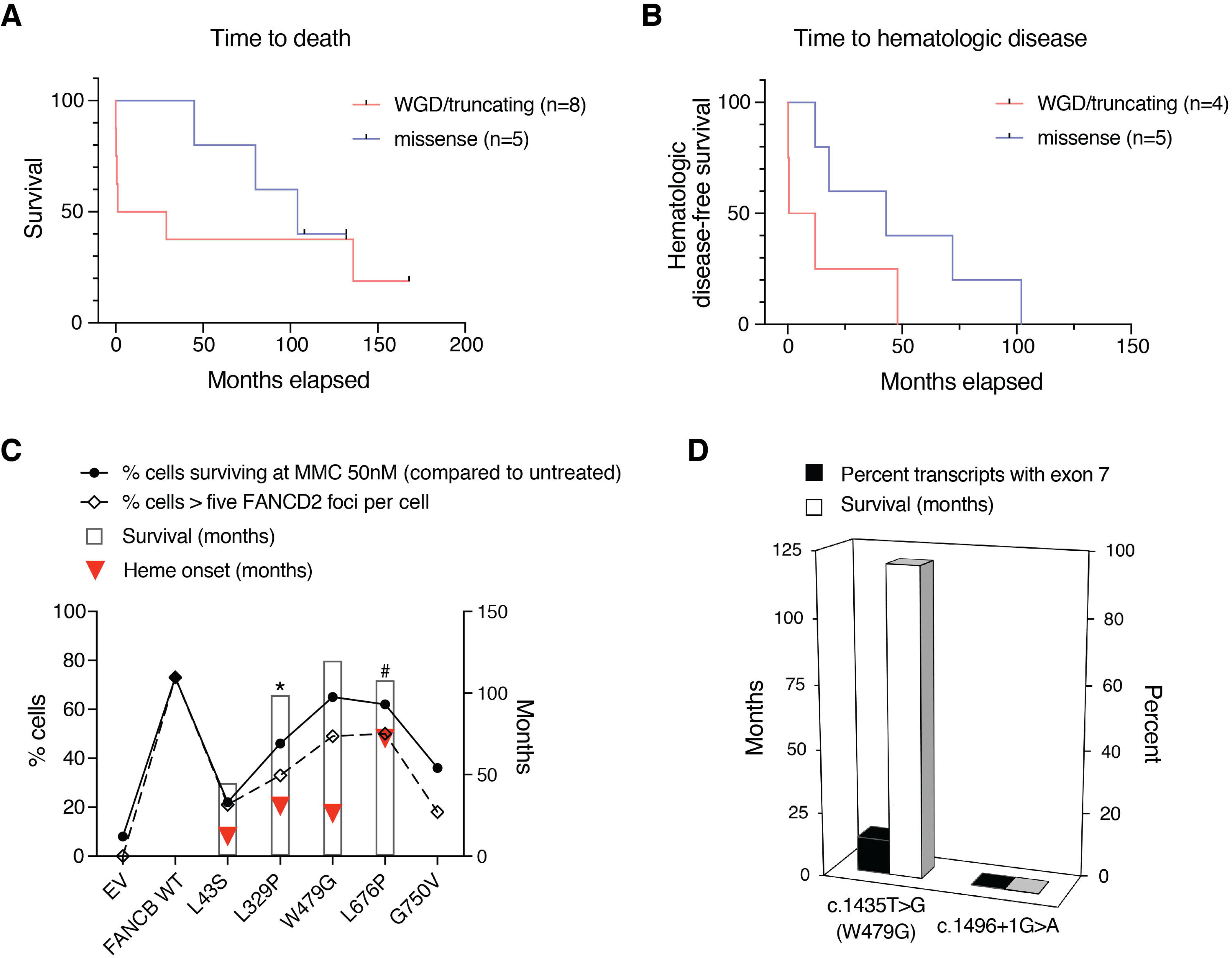
Clinical severity correlates with *in vitro* residual FANCB function. (A) The median time to death was earlier in the WGD/truncation variant group (Individuals 1, 2, 3, 4, 6, 11, 13 and 14) than in the missense group (individuals 16, 17, 18, 20 and 21) but was not statistically significant (Log-Rank test = 0.5198, p-value = 0.4709; HR = 1.693 (95% CI: 0.398 - 7.198)). Individuals excluded from this analyses are detailed in the supplemental methods. (B) The median time to the hematologic disease was earlier in the WGD/truncation variant group (individuals 3, 6, 11 and 14), than in the missense group (individuals 16, 17, 18, 20 and 21), but was also not statistically significant (Log-Rank test = 2.3962, p value = 0.1216; HR = 3.058 (95% CI: 0.669 - 13.973)). Individuals excluded from this analyses are detailed in the supplemental methods. (C) Survival and onset of hematologic manifestations (heme onset defined as the onset of bone marrow failure, MDS or leukemia, whichever comes first) tends to correlate with residual function, such as MMC sensitivity or quantity of FANCD2 foci. (D) The majority of the transcripts in the individual carrying c.1435T>G (predicted to express p.W479G) variant showed aberrant splicing, exon 7 skipping, which should result in an out-of-frame truncated protein. Though the expression level of c.1435T>G encoding the missense variant p.W479G is lower, its functional characteristics were near-normal, correlating with the relatively milder phenotype in this individual. In contrast, the individual carrying splice site variant (c.1496+1G>A), which resulted in the only product that lacked exon 7, passed away on postnatal day 1.

## DISCUSSION

### Individuals with FANCB pathogenic variants present with a wide spectrum of distinct, de novo or inherited variants

FANCB represents only ~2% of FA individuals, and we describe here 16 FANCB families, each with distinct variants (Figure 1, Table 1). Other previously reported FANCB variants were also only found in single families,^5,8,18,19,25-30^ suggesting an absence of founder mutations. This is likely due to the short lifespan of male carriers and the severity of symptoms, which probably extends to infertility as in Fancb^y/-^ mice.^31^ The nature of the variants included large deletion/duplication, missense, nonsense, splice junction, indel, and an unusual combination of overlapping *de novo* missense and indel variants. Deletions/duplications accounted for 20-30% of all variants in this and other studies.^5,8,27^ Deletions range from small (~3 kb, removing a single exon^5^) to large (520kb - 590kb, eliminating the entire *FANCB* gene along with flanking genes, individuals 1-3). Identification of such *FANCB* variants relies on methods for identifying large deletions such as aCGH.^32^ Confirming of the *de novo* status of *FANCB* variants, by eliminating the possibility of mosaicism in the parents, requires specialized techniques, such as deep sequencing, which is a robust method to detect unstable variants at low frequencies. In our cases, we found no evidence for mosaicism in probands with *de novo* and inherited variants, nor heterozygous carrier mothers/siblings (Table S2 and S5). However, for individual 15, who carried two overlapping *de novo* variants, we observed that the two variants existed at varying frequencies in peripheral blood DNA and the LCL. The reduced fraction of the c.2249G>T (G750V) variant in blood, and to an even greater extent, in LCL suggests that cells carrying indel variant transcript have a selective growth advantage. This could be due to the unstable nature of the missense variant that was observed both in patient LCL cells and in overexpressed FANCB null cells (Figure 3A and S2). These findings also suggest that the missense variant originated prior to the indel variant. Observations that transcripts differ from predictions based on genomic variants demonstrate the importance of including transcript analysis as a part of comprehensive molecular diagnosis.

**Table 1.**
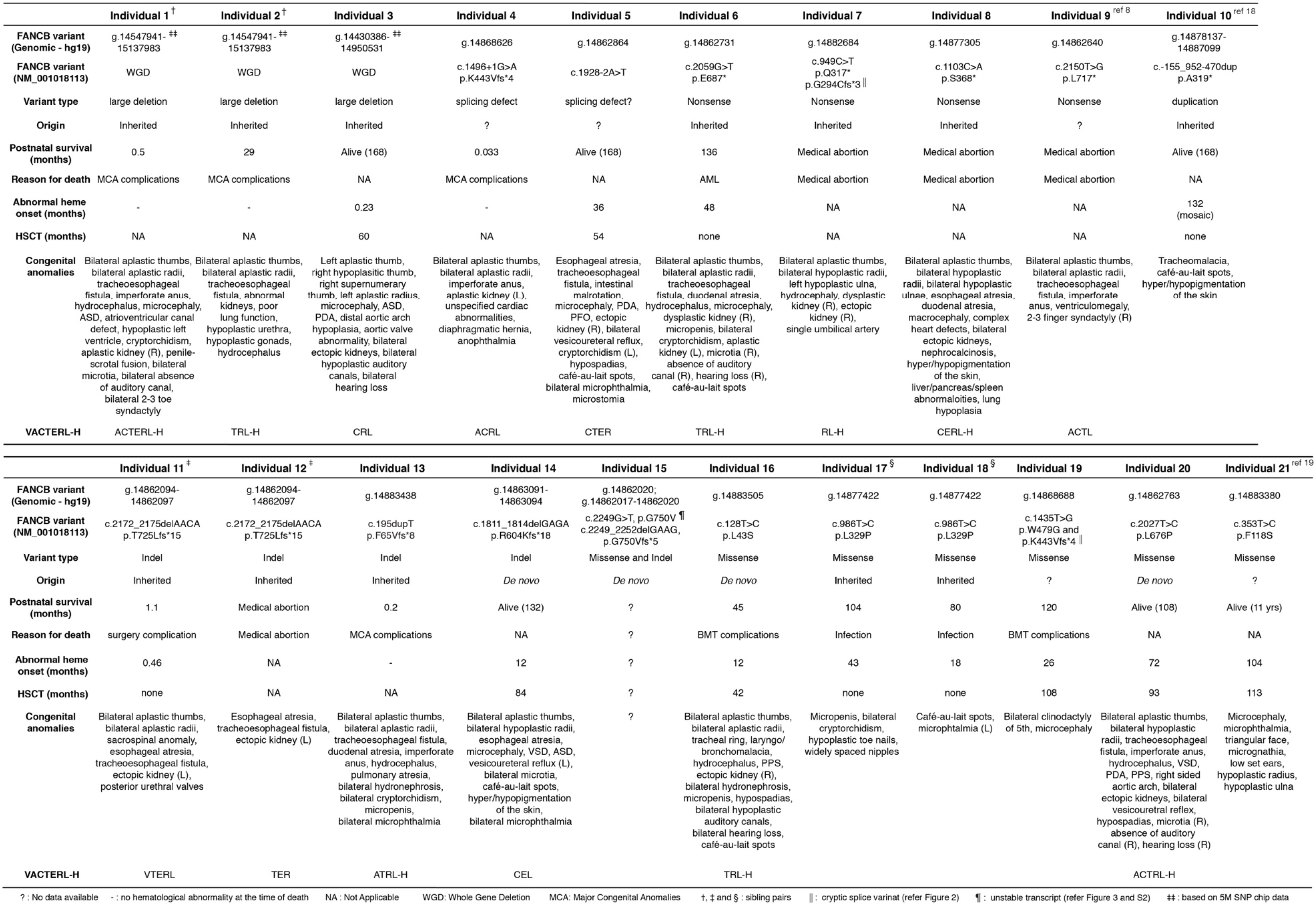
Molecular diagnosis and clinical presentations of individuals with *FANCB* pathogenic variants.

### Aberrant splicing from a nonsynonymous variant alters clinical course of the disease

Exon skipping played a role in our *FANCB* FA cohort, and the level of retained missense transcript (and hence full length FANCB protein) also correlated with clinical features. The exon 7 nonsynonymous variant, c.1435T>G (p.W479G), in individual 19 induces exon 7 skipping in about 85% of transcripts, reducing the contribution of the predicted nonsynonymous protein variant, p.W479G to <15% (Figure 2B-E). Functional studies revealed that the W479G FANCB protein retains substantial levels of FANCD2 ubiquitylation, foci formation, and ability to rescue null cell viability (Figure 3). In contrast, individual 4 with the splice donor variant c.1496+1G>A, generates the same aberrant transcript with exon 7 skipped, but in 100% of transcripts (Figure 2B-E). Individual 4 died 1 day after birth, whereas individual 19 lived until 120 months, dying of HSCT complications (Figure 6D). We, and others, have reported earlier that aberrant splicing is associated with certain synonymous, nonsynonymous, and nonsense variants in FA genes^27,15,33^ indicating that quantitative RNA analysis should be included to evaluate the molecular consequence of a genomic variant, particularly in the absence of clearly pathogenic alterations, but an apparent FA phenotype.

### Insight into the molecular function of *FANCB* from analysis of patient associated variants

Little is known about the actual function of FANCB protein within the FA core complex. Previous studies showed that the protein is stable for purification only as a recombinant heterodimer with FAAP100.^11^ Imaging of the FANCB:FAAP100 complex by EM suggests that the proteins form an intricately intertwined complex containing two of each subunit. We hypothesized that this unusual arrangement could place two FANCL molecules, each in close proximity to the ubiquitination sites of subunit of the FANCD2:FANCI heterodimeric substrate.^12^ Interestingly, we find that many variants in FANCB have a greater effect on FANCI monoubiquitination, and only mild or no effect on FANCD2 monoubiquitination *in vitro*. It is possible that FANCB, as the central dimerization mediator, coordinates the sequential monoubiquitination of FANCD2 and FANCI. Importantly, one of the longest reported truncations in FANCB completely lacks any detectable monoubiquitination activity, even though it retains association with FANCL, FAAP100 and the remainder of the FA core complex. This suggests that the C-terminus of FANCB has an essential role in the catalytic activity of the entire FA core complex. From this finding, we can conclude that even small FANCB truncations will be pathogenic.

We also identified two variants in the N-terminus of FANCB that cause an abrogation of interaction with the remainder of the FA core complex. The ~60-80% reduction in core complex association is sufficient to reduce the kinetics of FANCD2 and FANCI monoubiquitination by a similar rate. In a cellular context, where FA core complex concentrations are much more limiting (~0.1 ppm in Protein Abundance Database),^34^ such a reduced rate of association is likely to have an even greater effect on FANCD2 monoubiquitination. Together these findings suggest that a predicted beta-propellar in the N-terminus of FANCB mediates association with the FA core complex, while the remainder of the protein mediates the dimerization and catalytic activity of FANCL E3 RING ligase.

### Milder clinical presentation relates to the residual function of the FANCB missense variant

In this study, individuals with WGD/truncation variants in *FANCB* have more severe phenotypes than individuals with missense variants, although our analyses are limited by small sample sizes (Table 1, Figure 6 A-B). While it is ideal to perform the time to death analysis with the HSCT status as a time-varying covariate, we do not have a large enough sample size to have a meaningful conclusion from such analysis. Therefore, it was not reported. Results from all the functional assays on missense variants, both from cellular (Figure 3) and *in vitro* assays (Figure 4 and 5) as well as transcript analysis are summarized in Table 2. All of the missense variants tested were found to be pathogenic by one or more assay. In particular, compared to WT FANCB, the majority of variants showed relatively lower levels in the nucleus (Figure S4), which is also a characteristic of pathogenic *FANCA* missense variants.^33^ Together, the functional assays revealed a spectrum of activity, and the cell-based and protein-based assays broadly agreed on the extent of loss of function. The residual effect of each missense variant on cell viability and FANCD2 foci formation appeared to have some correlation with the hematological disease onset and overall survival of the individuals with *FANCB* missense variants (Figure 6C).

**Table 2.** Summary of functional assays for *FANCB* missense variants.

| FANCB variant | Transcript stability/<br>aberrant splicing<br>in patient cell line | Expression in null cells |  |  | Expression in insect cells |  | *In vitro* ubiquitination |  |
| --- | --- | --- | --- | --- | --- | --- | --- | --- |
|  |  | D2 Ub<br>upon MMC | % cell with<br>> 5 D2 foci<br>upon MMC | % cell<br>survival at<br>50 nM MMC | co-purification<br>with FANCL<br>and FAAP100 | % co-purification<br>with FANCA,<br>FANCC and FANCE | *K* for D2<br>mono-Ub<br>(%.min<sup>-1</sup>) | *K* for I<br>mono-Ub<br>(%.min<sup>-1</sup>) |
| WT | - | very strong | 73 | 75 | normal | normal | 18.47 | 1.1 |
| L43S | normal | very faint | 20 | 17 | normal | ~30-50 of WT | 6.09 | 0.09 |
| F118S | - | - | - | - | normal | ~20-40 of WT | 6.1 | 0.1 |
| L329P | normal | faint | 32 | 45 | normal | normal | 11.51 | 0.09 |
| W479G | exon7 skipped in 86.5% | strong | 48 | 69 | - | - | - | - |
| L676P | normal | strong | 49 | 61 | normal | normal | 21.28 | 0.15 |
| G750V | unstable transcript | faint | 17 | 39 | normal | normal | 21.18 | 0.86 |
| L717* | - | - | - | - | normal | normal | 0.93 | 0 |

D2 - FANCD2, I - FANCI, Ub - Ubiquitination, *K* - rate constant

Because *FANCB* is X-linked and, to date, restricted to male cases, each *FANCB* patient carries only one deleterious allele and a generally more severe form of FA. Our study highlights how complementary approaches to variant analysis converge to provide meaningful new knowledge of both FANCB protein function, and the etiology of FANCB-associated FA.

## Supporting information

All supplemental data

## ACKNOWLEDGEMENTS

We thank the individuals and the families who participated in this study. We thank K.J Patel for the FANCE antibody. This work was supported by grants from the National Heart Lung and Blood Institute (R01 HL120922) (A.S.), National Cancer Institute (R01 CA204127) (A.S.), National Center for Advancing Translational Sciences (UL1 TR001866) (M.J., C.S.J., R.V., and A.S.), American Society of Hematology Scholar Award (M.J.), the National Health and Medical Research Council, Australia (APP1123100 and APP1145391) (A.J.D.) and the Victoria government IOS program (A.J.D.). A.S. is a HHMI Faculty Scholar. A.J.D. is a Victorian Cancer Agency mid-career fellow. R.R-B., F.X.D., D.C.K. and S.C.C. acknowledge the support from the Intramural Research Program of the National Human Genome Research Institute, NIH.

## CONTRIBUTION

M.J., R.R-B., S.v.T., F.P.L. V.M. and W.T. performed experiments. M.J., R.R-B., S.v.T., F.X.D., S.C.C. and A.J.D. analyzed results. R.R-B, D.C.K., F.X.D. and S.C.C. collected and analyzed the genomic sequence. M.J., R.R-B., and A.J.D. made the figures. R.O.R., F.P.L., and A.S. managed the IFAR. A.D.A. established the IFAR, obtained follow up clinical information and edited the paper. C.S.J. and R.V performed statistical analysis of clinical data. M.J., R.R-B., A.J.D., A.S. and S.C.C. designed the research and wrote the paper. R.O.R., P.M., F.P. and C.D. collected clinical presentations of some of the individuals.

