## Supplementary material for "Clinical severity in Fanconi anemia correlates with residual function of FANCB missense variants": All supplemental data

### Supplemental Methods

*Identification of disease-causing FANCB mutations*

Nextgen sequencing targeted and captured the entire genomic regions of all FA genes, including *FANCB*, using the TruSeq custom amplicon kit (Illumina), and sequenced using 101bp paired-end reads to a depth of ~500x using the HiSeq 2000 system (Illumina). Illumina Bead chips (SNP arrays), OmniExpressExome (1M SNPs) and Omni5exome (5.25M SNPs) were employed for mapping the interval of large *FANCB* deletions. Data collection from bead chips and analysis for deletion intervals were as described earlier.[1]

*Reverse transcription-PCR (RT-PCR)*

Screening for aberrant *FANCB* splice products was done using gene specific primers and KAPA2G Fast HotStart ReadyMix PCR Kit (Kapa Biosystems). The RT-PCR products were either sequenced directly, or after cloning (if multiple bands). qPCR was performed using Platinum™ SYBR™ Green qPCR SuperMix-UDG (Thermo Fisher Scientific) and QuantStudio 12K-flex (Life Technologies). ΔΔCt was calculated and normalized to empty vector (EV).

*Amino acid sequence conservation*

The Consurf alignment sequence logo was derived using consurf analysis (http://consurf.tau.ac.il/2016/) of human FANCB and 98 homologs with sufficient coverage for the analysis.[2] The output alignment was analyzed in Geneious software (Biomatters development team) to generate the sequence logos.

*Individual specific qf-PCR*

To measure relative levels of *FANCB* transcripts with or without exon 7 in individual 4 and 19, two forward primers, M13F-tailed ex7-8 junction FP and M13F-tailed ex6-8 junction FP, that bind to the exon 7-8 junction and the exon 6-8 junction, respectively, and a reverse primer PIG-tailed ex8 RP were used. The expected product from transcript with exon 7 is 3 bp longer than that from without exon 7. Average peak amplitude from eight qf-PCR reactions was taken to calculate average percent splice products.

In individual 15, the relative levels of *FANCB* missense and indel variant at genomic DNA was measured using a forward and reverse primer, M13F-tailed int9 FP and PIG-tailed ex10 RP, respectively. Relative transcript levels were measured using M13F-tailed ex9 FP and PIG-tailed ex10 RP primers. The expected product with indel variant (c.2249_2252delGAAG) is 4 bp smaller than the product with missense variant (c.2249G>T). The average peak amplitude from four independent experiments was taken to calculate average percent of missense and indel variants.

*MiSeq sequencing*

MiSeq sequencing was performed according to Illumina protocol. The primers are listed in the Table S2. MiSeq sequence reads were quantified using samtools.[3] PCR products that included individual-specific variants were generated using KAPA2G Fast HotStart ReadyMix (KAPA Biosystems) and 2 uM HPLC purified primers. A subsequent PCR reaction was performed with the first reaction as the template, and adapter primers with sample-specific barcodes. Cluster generation and sequencing of PCR products were performed with the MiSeq Reagent Kit v3 with 2x300 paired-end reads on an Illumina MiSeq instrument, giving an output of ~10 million total reads (Illumina).

*MiSeq data analysis using Sam-tools*

For a given variant, the following command was implemented: *> samtools view -q 30 -f 2 input.bam I grep 'sequence' I wc -l*, where "input.bam" is the sample .bam file, and "sequence" refers to the *FANCB* variant position plus at least 5 bps upstream and downstream of the variant's position. Quantification was performed for the wild-type sequence as well as the variant sequence. The output, or read count "wc -l", for the wild-type and variant sequences were summed and used as the denominator to calculate percentage reads. For each *FANCB* SNV, in addition to the variant and wild-type sequences, the number of reads were counted for each nucleotide at the position of the variant, that was neither wild-type nor the initially queried variant, to gauge the level of expected background sequence reads.

*Complementation of FANCB deficient cell lines with wild type (WT) or mutant FANCB*

The WT *FANCB* and missense variants (generated using QuikChange II site-directed mutagenesis, Agilent Technologies) were subcloned into pDONR223 vector, subsequently recombined into lentiviral vector with pHAGE CMV N-terminus HA-FLAG as described previously.[19] The lentiviral plasmids were packaged into lentivirus using Trans-IT (Mirus Bio) transfection of HEK293T cells. FANCB-null fibroblasts (RA2275) or selected individual lines were infected with lentivirus carrying empty vector (EV) or WT or missense-variant-expressing *FANCB* cDNA and selected by puromycin. Western blotting of FANCD2 (NB100-182, Novus), HA-FANCB (16B12, BioLegend) and $\alpha$-Tubulin (DMA1, Sigma Aldrich) and immunofluorescence of FANCD2 and HA-FANCB foci was performed with and without 24hr exposure to 1$\mu$M mitomycin C (MMC) as previously described.[19,20] Cell survival was measured after increasing doses of MMC treatment using Z2 Coulter® Particle and Size Analyzer (Beckman Coulter).

*Extraction of nuclear proteins*

Nuclear extraction was performed as previously described with some modifications.[4] Cells were harvested with 0.25% Trypsin-EDTA (Gibco) and washed with Dulbecco's Phosphate-Buffered Saline. Buffer A, which is consisted of 10mM HEPES (Sigma Aldrich), 1.5mM MgCl2 (Sigma Aldrich), 10mM KCL (Fisher Chemical), 0.5mM DTT (Sigma Aldrich), 0.05% Nonidet-P40 (US Biological) at pH 7.9, supplemented with 0.3M PMSF (Thermo Fisher Scientific) and cOmplete™, mini, EDTA-free protease inhibitor cocktail (Roche) immediately before use, was added to cell pellets and mixed and cells were kept on ice for 10 minutes. Samples were centrifuged at 4 °C at 3000 RPM for 10 minutes. Supernatant containing cytoplasmic fraction was aspirated and nuclear pellets were resuspended in 100uL of 2X Laemmli sample buffer (Bio-Rad) supplemented with 5% 2-Mercaptoethanol (Sigma Aldrich), boiled at 100C for 10 minutes, and used for western blotting of HA-FANCB, PCNA (PC-10, Santa Cruz Biotechnology) and $\alpha$-Tubulin.

*Recombinant protein purification & expression*

For the BL100 construct, N-terminal Flag-tagged FANCB was cloned into pFL-EGFP.[5] The codon optimized pFL-EGFP FANCBopt was used as a template to make FANCB F118S, L676P, L717X, G750V by mutagenesis, or ordered as a Gene Block (L43S, L329P). pFL-EGFP FANCB constructs were combined with pSPL FAAP100 and FANCL (isoform 2) by Cre recombinase, selected on Ampicillin and Spectinomycin, confirmed by restriction digest and sequencing, before insertion into Multibac bacmid via TN7 transposition.[6]

Recombinant BL100 WT and mutant proteins were produced from High Five cells (Invitrogen) infected with Multibac baculoviruses at MOI of 2. Cell pellets were washed with 1x PBS and re-suspended in 3 pellet volumes buffer A (20mM TEA, pH7.5, 200mM NaCl, 10% glycerol, 1mM DTT, protease inhibitors) (Sigma P8340). Lysates were cleared by centrifugation at 50,000 x g for 30min. 1ml M2 Flag resin (Sigma A2220-10ML) was added to lysates and incubated for 1 hour with gentle mixing. Resin was washed 4x and eluted in Buffer A (omitted protease inhibitors) + 100ug/ml Flag peptide. All purifications done at 4°C.

*Sub-complex assembly with BL100 WT or Mutants*

Interaction of BL100 WT or mutant proteins with recombinant FANCA-FANCG-FAAP20 (AG20) and FANCC-FANCE-FANCF (CEF) was done by expressing each sub-complex separately, or by co-infection. After washing with 1x PBS, pellets were re-suspended in buffer A, and equal volumes of CEF and AG20 lysate were added to BL100 WT or mutant lysates. Once the lysates were mixed, they were incubated at 4C for 30min with gentle agitation. Then lysates were cleared by centrifugation, and Flag affinity tag purified as described above for BL100. The following antibodies were used to verify protein interactions: rabbit polyclonal antibodies against FANCA and FANCG (Fanconi Anemia Research Fund), FANCL (GTX100162, GeneTex), FANCB (D9W6S, Cell Signaling), FANCE,[7] and FAAP20 and FAAP100 (Sigma).

*In vitro ubiquitination experiments*

Standard ubiquitination reactions contained 10mM recombinant human HA-ubiquitin (Boston Biochem U-110-01M), 50 nM human recombinant Ube1(CSIRO), 100 nM FANCT/UBE2T, 2 mM adenosine triphosphate in reaction buffer (50 mM Tris [pH 7.4], 2.5 mM MgCl2, 150 mM NaCl, 0.01%Tritonx100, and 1 mM DTT). Each reaction contained 165 nM FANCI:D2, 165 nM BL100 WT or mutant protein and 165 nM CEF. FANC ID2 and CEF purifications as described in van Twest et al.[8] Reactions were set up on ice, incubated at 25°C for 30min unless otherwise specified. All reactions were stopped by adding 10mM NuPage LDS sample buffer (Invitrogen), and heated at 80°C for 5min. Samples were separated by SDS-PAGE using BOLT 4%–12% Bis-Tris or NuPAGE 3%–8% Tris-Acetate gels (Invitrogen). For ubiquitination western blots, FANCD2 was detected with StrepII tag antibody (ab76949, Abcam), and FANCI was detected with Flag antibody (OAEA00002, Aviva Systems Biology). Quantification was performed as per van Twest et al.[8]

*Statistical analyses of clinical data*

Time to death and time to hematologic disease (defined as the time at which any of the following events: hemoglobin < 10 g/dL, platelets < 100 K/uL, absolute neutrophil count < 1000/uL, myelodysplastic syndrome or leukemia) were analyzed by Kaplan-Meier method to determine the median time to event depending on mutation group (i.e. whole gene deletion(WGD)/truncation vs. missense). The Log-Rank test was used to test the homogeneity of the survival curves, and the corresponding hazard ratios and 95% confidence intervals were computed from a Proportional Hazards regression model. Analyses were performed using SAS Studio Version 3.7 and GraphPad v8.0.1.

Among 21 individuals included in this study (19 from IFAR and 2 from reported studies[9,10]), we excluded two individuals from both time to death and time to hematologic disease analyses due to known somatic mosaicism (individual 10) and a lack of clinical information (individual 15). Four medical abortions were also excluded from the both analyses (individuals 7, 8, 9 and 12). Additional two individuals were excluded specifically for mutation-based (WGD/truncation vs. missense) analyses because of the presence of both truncating and missense variants (individual 19) and a lack of RNA data to confirm the consequence at protein level from a splice junction variant (individual 5). Another four individuals were excluded only from time to hematologic disease analysis due to a lack of disease onset information (individuals 1, 2, 4 and 13). The final analysis included 13 individuals for time to death and nine individuals for time to hematologic disease.

### Supplemental Tables

**Table S1. qf-PCR primers used in the study**

| Primer | Sequence |
|---|---|
| FAB M13F-tailed ex7-8 junction FP | tgtaaaacgacggccagtACATCTTCTTTGAAGCTGTCC |
| FAB M13F-tailed ex6-8 junction FP | tgtaaaacgacggccagtCAAGTGCAGAGGAGGTCC |
| FAB PIG-tailed ex8 RP | gtgtcttCTGCCACACACAACATAACG |
| M13F-FAM | /6-FAM/TGTAAAACGACGGCCAGT |
| βactin M13F-tailed FP | tgtaaaacgacggccagtCACCAACTGGGACGACAT |
| βactin PIG-tailed FP | gtgtcttACAGCCTGGATAGCAACG |
| FAB M13F-tailed ex9 FP | tgtaaaacgacggccagtAGACCGGGAAGTTTCTATGG |
| FAB M13F-tailed int9 FP | tgtaaaacgacggccagtTGACTGCTTTGTGCTGTCCT |
| FAB PIG-tailed ex10 RP | gtgtcttCCTTTGCTCACTTCACACCT |

**Table S2. MiSeq data for *de novo* variants**

| Individual 15 | | | c.2249G>T; c.2249_2252delGAAG | | | | | | | |
|---|---|---|---|---|---|---|---|---|---|---|
| | | | Proband - PB | | Proband - LCL | | Father - PB | | Mother - PB | |
| WT | TAAAATCAGGAAGTGAGAATT | | 2169 | 0.92% | 1396 | 0.58% | 184015 | 99.76% | 222366 | 99.90% |
| c.2249G>T | TAAAATCAGTAAGTGAGAATT | | 188348 | 80.13% | 128029 | 53.08% | 430 | 0.23% | 221 | 0.10% |
| c.2249_2252delGAAG | TAAAATCAG- - - -TGAGAATT | | 44543 | 18.95% | 111774 | 46.34% | 13 | 0.01% | 2 | 0.00% |
| | | Total Reads | 235060 | | 241199 | | 184458 | | 222589 | |
| **Individual 16** | | | **c.128T>C** | | | | | | | |
| | | | **Proband - PB** | | **Proband - LCL** | | **Father - PB** | | **Mother - PB** | |
| WT | ACACCCATATTACATGTCAG | | 3151 | 1.05% | 3135 | 0.86% | 413100 | 99.57% | 409011 | 99.55% |
| c.128T>C | ACACCCATATCACATGTCAG | | 296850 | 98.95% | 360540 | 99.14% | 1765 | 0.43% | 1834 | 0.45% |
| | | Total Reads | 300001 | | 363675 | | 414865 | | 410845 | |
| **Individual 14** | | | **c.1811_1814delGAGA** | | | | | | | |
| | | | **Proband - PB** | | **Proband - LCL** | | **Father - PB** | | **Mother - PB** | |
| WT | CAAATTATGGAGAGAGAAAGTGGTAACTG | | 168 | 0.04% | 107 | 0.02% | 456112 | 100.00% | 608601 | 100.00% |
| c.1811_1814delGAGA | CAAATTATGGA- - - -GAAAGTGGTAACTG | | 440114 | 99.96% | 453456 | 99.98% | 17 | 0.00% | 11 | 0.00% |
| | | Total Reads | 440282 | | 453563 | | 456129 | | 608612 | |
| **Individual 20** | | | **c.2027T>C** | | | | | | | |
| | | | **Proband - PB** | | **Proband - Fib** | | **Father - PB** | | **Mother - PB** | |
| WT | AGGTGTGGCTCTTAGAACATATG | | 1428 | 0.52% | 1482 | 0.55% | 282225 | 99.18% | 311130 | 99.04% |
| c.2027T>C | AGGTGTGGCCCTTAGAACATATG | | 272896 | 99.48% | 269966 | 99.45% | 2344 | 0.82% | 3001 | 0.96% |
| | | Total Reads | 274324 | | 271448 | | 284569 | | 314131 | |
| PB - Peripheral Blood; LCL - Lymphoblastoid Cell Lines; Fib - Fibroblast | | | | | | | | | | |

**Table S3. ConSurf score for missense variants**

| Missense variant | ConSurf score | % Frequency human aa in 99 FANCB homologs | Frequency variant aa in 99 FANCB homologs |
|---|---|---|---|
| L43S | 9 | 97 | 0 |
| F118S | 7 | 94 | 0 |
| L329P | 8 | 96 | 0 |
| W479G | 9 | 100 | 0 |
| L676P | 8 | 96 | 0 |
| G750V | 4 | 81 | 0 |

### Table S4. MiSeq primers used in the study

**First-round PCR primers**

| Samples | FWD Primer Name | Adapter Seq + *gene specific FWD primer seq* | REV Primer Name | Adapter Seq + *gene specific REV primer seq* | size | Size with adapters |
|---|---|---|---|---|---|---|
| Individual 13 and 16 | FANCB_ex.3b_F | ACACTCTTTCCCTACACGACGCTCTTCCGATCT*GGCCATCCTTCATCTCATAGC* | FANCB_ex.3b_R | GTGACTGGAGTTCAGACGTGTGCTCTTCCGATCT*TTTAGAAACATCAACTGGAATCAAT* | 469 | 605 |
| Individual 7 | FANCB_ex.3d_F | ACACTCTTTCCCTACACGACGCTCTTCCGATCT*TTAGACAATAAGACTCCAGAATGAACT* | FANCB_ex.3d_R | GTGACTGGAGTTCAGACGTGTGCTCTTCCGATCT*CTGAGGAAGAATGTACTCAAGAGC* | 474 | 610 |
| Individual 8, 17 and 18 | FANCB_ex.4_F | ACACTCTTTCCCTACACGACGCTCTTCCGATCT*TTTGTTGAAAGGAGATTCAATG* | FANCB_ex.4_R | GTGACTGGAGTTCAGACGTGTGCTCTTCCGATCT*TGAGTTTTACAAATGACAACTACATGA* | 432 | 568 |
| Individual 14 | FANCB_ex.8c_F | ACACTCTTTCCCTACACGACGCTCTTCCGATCT*TCATTAAACTCTGCCCATTATCA* | FANCB_ex.8c_R | GTGACTGGAGTTCAGACGTGTGCTCTTCCGATCT*CTTTGTTTGTGAACACCCATCT* | 330 | 466 |
| Individual 20 | FANCB_ex.9_F | ACACTCTTTCCCTACACGACGCTCTTCCGATCT*GGTAATTTTGTTGGCACTTTTAG* | FANCB_ex.9_R | GTGACTGGAGTTCAGACGTGTGCTCTTCCGATCT*CTTCCCGGTCTTTCACAAAA* | 285 | 421 |
| Individual 11, 12 and 15 | FANCB_ex.10a_F | ACACTCTTTCCCTACACGACGCTCTTCCGATCT*CCTCTGCATAAAATTGCTTTCA* | FANCB_ex.10a_R | GTGACTGGAGTTCAGACGTGTGCTCTTCCGATCT*TCCCGTTACCTCTCCTCCAC* | 487 | 623 |

**Second-round PCR primers**

| | Adapter Seq + Index Seq (*) + *Overlap Seq* |
|---|---|
| **FWD Primer** | AATGATACGGCGACCACCGAGATCTACAC********_ACACTCTTTCCCTACAC_ |
| **REV Primer** | CAAGCAGAAGACGGCATACGAGAT********_GTGACTGGAGTTCAG_ |

### Table S5. MiSeq data for inherited variants

| | | | | | | | |
|---|---|---|---|---|---|---|---|
| **Individual 11** | | **c.2172_2175delAACA** | | | | | |
| | | **Proband - PB** | | **Mother - PB** | | **Mother - LCL** | |
| WT | TTTAGGAATCAAACAGTTATGTTCC | 94 | 0.05% | 99245 | 45.74% | 117841 | 53.65% |
| c.2172_2175delAACA | TTTAGGAATCA- - - -GTTATGTTCC | 193697 | 99.95% | 117748 | 54.26% | 101805 | 46.35% |
| | Total Reads | 193791 | | 216993 | | 219646 | |
| **Individual 18** | | **c.986T>C** | | | | | |
| | | **Proband - PB** | | **Unaffected Sibling - PB** | | | |
| WT | TTAGTACTGATAGATGA | 4650 | 0.81% | 234082 | 47.85% | | |
| c.986T>C | TTAGTACCGATAGATGA | 566791 | 99.19% | 255130 | 52.15% | | |
| | Total Reads | 571441 | | 489212 | | | |
| background 1 | TTAGTACGGATAGATGA | 242 | | 491 | | | |
| background 2 | TTAGTACAGATAGATGA | 1178 | | 646 | | | |
| **Individual 13** | | **c.195dupT** | | | | | |
| | | **Proband - PB** | | **Father - PB** | | **Mother - PB** | |
| WT | ATTTTTTACCATAAAGGAAG | 7837 | 1.89% | 385372 | 99.93% | 188039 | 54.12% |
| c.195dupT | ATTTTTTTACCATAAAGGAAG | 406122 | 98.11% | 270 | 0.07% | 159409 | 45.88% |
| | Initial Total Reads | 363621 | | | | 143069 | |
| | Additional reads with 'A' mismatch a | 42499 | | | | 16340 | |
| | Total Reads | 413959 | | 385642 | | 347448 | |
| background 1 | ATTTTTTAACCATAAAGGAAG | 27 | | 24 | | 26 | |
| background 2 | ATTTTTTGACCATAAAGGAAG | 20 | | 19 | | 22 | |
| background 3 | ATTTTTTCACCATAAAGGAAG | 17 | | 16 | | 20 | |
| **Individual 7** | | **c.949C>T** | | | | | |
| | | **Proband - Fib** | | **Father - PB** | | **Mother - PB** | |
| WT | AGAGCTTTCAGGTACAACA | 784 | 0.36% | 232207 | 99.89% | 125234 | 57.03% |
| c.949C>T | AGAGCTTTTAGGTACAACA | 218462 | 99.64% | 266 | 0.11% | 94353 | 42.97% |
| | Total Reads | 219246 | | 232473 | | 219587 | |
| background 1 | AGAGCTTTGAGGTACAACA | 24 | | 35 | | 90 | |
| background 2 | AGAGCTTTAAGGTACAACA | 137 | | 105 | | 52 | |
| **Individual 8** | | **c.1103C>A** | | | | | |
| | | **Proband - Fib** | | **Father - PB** | | **Mother - PB** | |
| WT | AAATAAACTATTCGGTAAGTTCTATT | 4516 | 1.09% | 409630 | 99.36% | 220265 | 51.95% |
| c.1103C>A | AAATAAACTATTAGGTAAGTTCTATT | 408533 | 98.91% | 2657 | 0.64% | 203689 | 48.05% |
| | Total Reads | 413049 | | 412287 | | 423954 | |
| background 1 | AAATAAACTATTTGGTAAGTTCTATT | 220 | | 689 | | 456 | |
| background 2 | AAATAAACTATTGGGTAAGTTCTATT | 1224 | | 63 | | 481 | |

PB - Peripheral Blood; LCL - Lymphoblastoid Cell Lines; Fib - Fibroblast

### Supplemental Figures

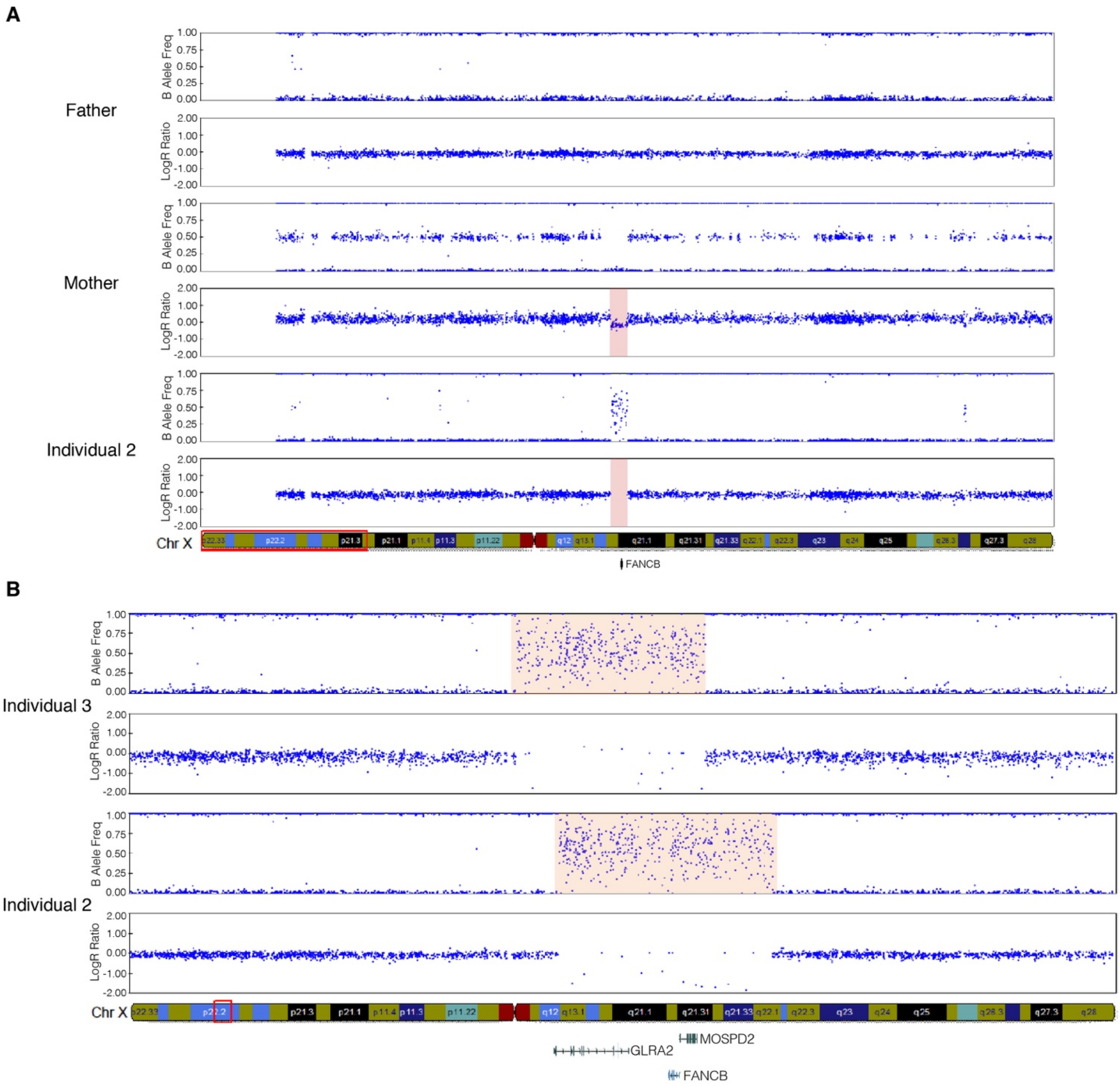

**Figure S1. Large deletions in two FANCB families.**

(A) SNP genotype data from individual 2 trio using OmniExpressExome (1M SNPs) arrays shows a hemizygous *FANCB* whole gene deletion in the proband and heterozygous deletion in the mother.

(B) High-density SNP data showing *FANCB* whole gene deletion intervals from individuals 2 and 3. The deletion coordinates, from HumanOmni5QuadExome_v1.2 (5.25 M SNPs), for individual 2 and 3 are hg19_chrX:14547941-15137983 (590 kb) and hg19_chrX:14430386-14950531 (520 kb), respectively.

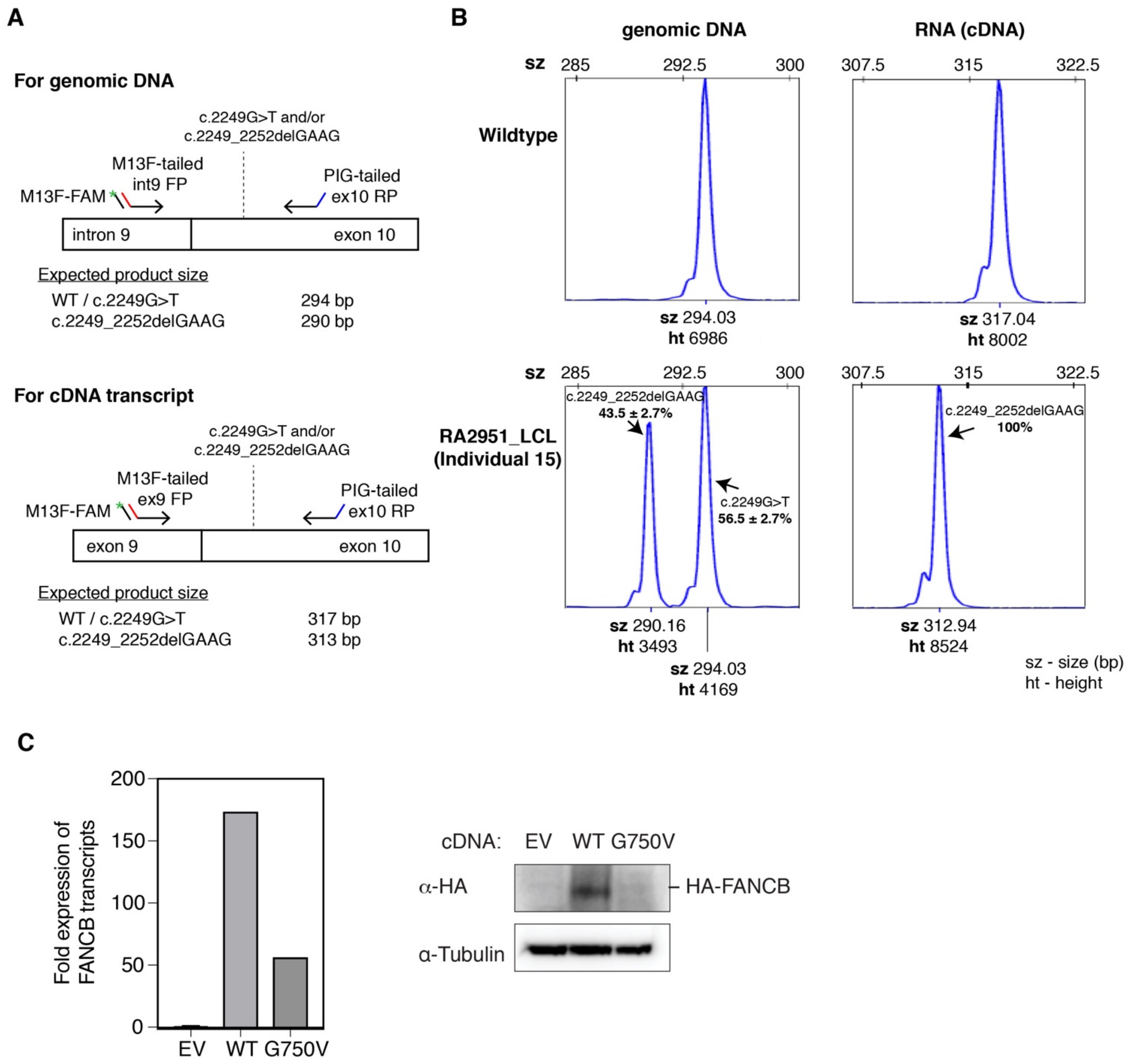

**Figure S2. Relative quantification of two variants in individual 15 at genomic DNA and transcript level.**

(A) Schematics of quantitative fluorescence PCR (qf-PCR) to measure relative levels of c.2249G>T and c.2249_2252delGAAG variant in individual 15. For genomic DNA, a *FANCB* specific forward primer in intron 9 (M13F-tailed int9 FP), and a reverse primer in exon 10 (PIG-tailed ex10 RP) were used to amplify products. The expected product size with c.2249G>T and c.2249_2252delGAAG variant is 294bp and 290bp, respectively. For cDNA transcript, a *FANCB* specific forward primer in exon 9 (M13F-tailed ex9 FP), and same reverse primer (PIG-tailed ex10 RP) were used to amplify products. The expected product size with c.2249G>T and c.2249_2252delGAAG variant is 317bp and 313bp, respectively. Corresponding sample from an unaffected individual was used as a control. FAM-labelled M13 forward primer (M13F-FAM; with green star) was included to generate fluorescently labelled products.

(B) Example qf-PCR peak profiles from ABI sequencer. Fragment size scales are shown at the bottom, and vertical scale marks the intensity of peaks. The fragment size (sz) and peak intensity (ht) are shown underneath each peak. The fragment size for products from both genomic DNA and cDNA transcript appeared around expected product sizes. The average peak amplitude from four independent reactions that correspond to c.2249G>T and c.2249_2252delGAAG variant was used to calculate their relative percentages.

(C) FANCB-null fibroblasts (RA2275) expressing EV or full-length FANCB WT or G750V missense variant cDNA were used for RT-qPCR and western blot. RT-qPCR was performed in three technical replicates. $\Delta\Delta$Ct was calculated for FANCB gene expression and normalized to that of EV. GAPDH was used as a reference gene. FANCB G750V transcripts were expressed less than one third of FANCB WT transcripts.

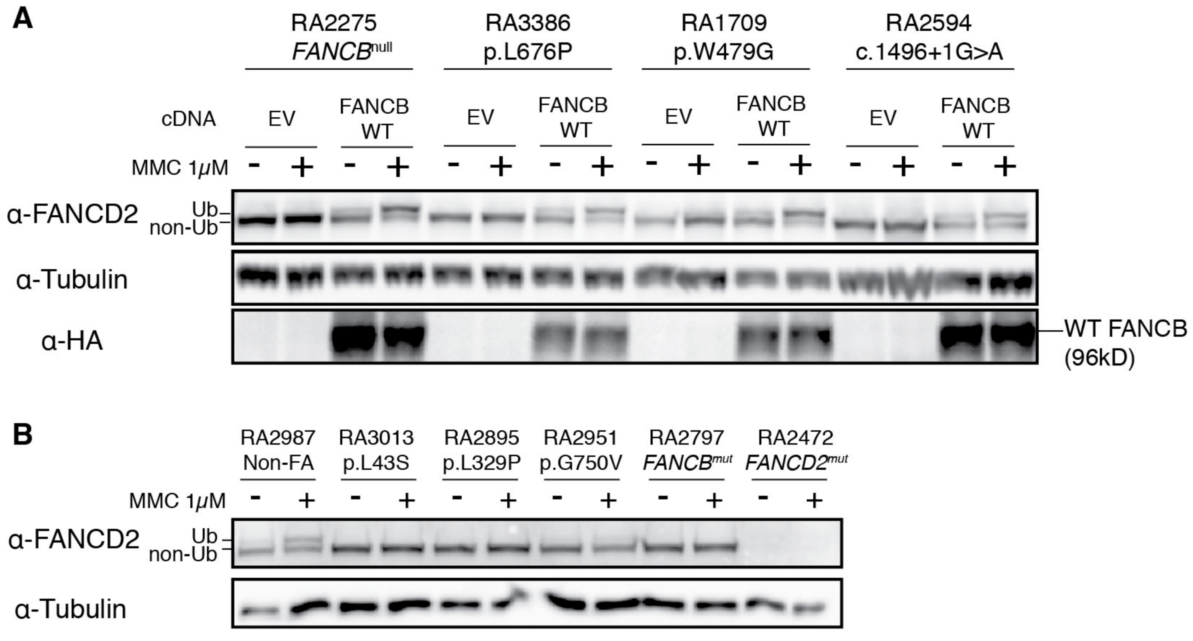

**Figure S3. Examination of FANCD2 ubiquitination upon MMC exposure in individual cell lines.** (A) Individuals fibroblasts carrying missense variants or splicing variant (c.1496+1G>A) were complemented with either empty vector (EV) or HA-FANCB WT cDNA, and then used for FANCD2 western blot with and without MMC exposure. All cell lines tested recovered FANCD2 ubiquitination by expression of HA-FANCB WT cDNA, confirming FANCB being the mutant gene causing FA in these individuals.

(B) FANCD2 western blot with and without MMC treatment in patient-derived LCL cell lines. RA2951 shows low level of FANCD2 ubiquitination upon MMC treatment. Note: Deep sequencing of RA2951 genomic DNA showed 53% of reads with p.G750V (c.2249G>T) variant and remaining reads with p.G750Vfs*5 (c.2249_2252delGAAG) variant. RNA analysis from RA2951 showed transcripts with only the indel variant (Figure S2B).

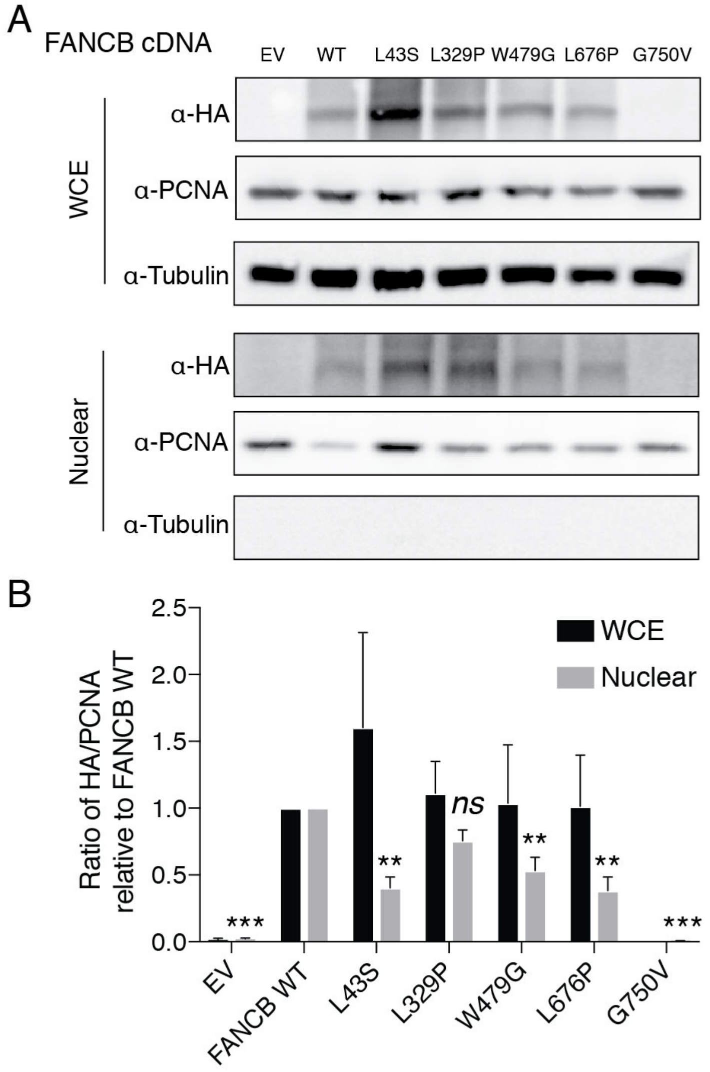

**Figure S4. Decreased nuclear localization of HA-FANCB missense variants.**

(A) Western blot of HA-FANCB in whole cell extract (WCE) and nuclear fraction (N).

(B) Relative intensity of bands was measured by ImageJ2 and normalized to that of FANCB WT. Statistical analysis was performed using one-way ANOVA followed by Dunnett's multiple comparison test. ** $P < 0.01$ and *** $P < 0.005$. *ns*: not significant.

### Structural Motifs Using Random Fields (SMURF)

```
>FANCB
Raw score: 51.4569
P-value: 2.227631e-04

--------------------------------------atvvtetvy-repaidsqgraf
                                         |||||| |||BBB|||BBB
MTSKQAMSSNEQERLLCYNGEVLVFQLSKGNFADKEPT---KTPILHVRRMVFDRGTKVF

atssngkillldltgskkvlfrpddnpspaviyd-----ntgkllvvtrp-----tnglv
BBB| |BBBBBB  |||BB|||    ||BBBB||     |||BBBBBB||     ||BBB
VQKS-TGFFTIK--EENSHLKI----MCCNCVSDFRTGINLPYIVIEKNKKNNVFEYFLL

iytlkwekktdiga-ialdnvndgafdtl-------kgalyvtdrkvgsvyifyldsgkv
BBB||||||||||| |||||||||BB|||       ||BBBBBB||||BBBBBB   |BB
ILHSTNKFEMRLSFKLGYEMKDGLRVLNGPLILWRHVKAFFFISSQTGKVVSVS---GNF

--tplfkglsingiaispdgltlyiret-------------trnkiwkfelgtslvlkda
  |||||||||||BBBB|| |BBBBBB|             |||BBBBBB||||||||||
SSIQWAGEIENLGMVLLGL-KECCLSEEECTQEPSKSDYAIWNTKFCVYSLESQEVLSDI

-------------tgpvgisvds-dgtlyvas--grsniaryklskdivqiesp-gksty
             |||||BBBB| ||BBBBBB  |||BBBBBB|||||BB|||| |||||
YIIPPAYSSVVTYVHICATEIIKNQLRISLIALTRKNQLISFQNGTPKNVCQLPFGDPCA

savspf---dgsvlves-snsslyilnwe-------------------------------
BBBB||   ||BBBBBB |||BBBBBB||
VQLMDSGGGNLFFVVSFISNNACAVWKESFQVAAKWEKLSLVLIDDFIGSGTEQVLLLFK

------------------------------------------------------------

DSLNSDCLTSFKITDLGKINYSSEPSDCNEDDLFEDKQENRYLVVPPLETGLKVCFSSFR

------------------------------------------------------------

ELRQHLLLKEKIISKSYKALINLVQGKDDNTSSAEEKECLVPLCGEEENSVHILDEKLSD

------------------------------------------------------------

NFQDSEQLVEKIWYRVIDDSLVVGVKTTSSLKLSLNDVTLSLLMDQAHDSRFRLLKCQNR

------------------------------------------------------------

VIKLSTNPFPAPYLMPCEIGLEAKRVTLTPDSKKEESFVCEHPSKKECVQIITAVTSLSP

------------------------------------------------------------

LLTFSKFCCTVLLQIMERESGNCPKDRYVVCGRVFLSLEDLSTGKYLLTFPKKKPIEHME

------------------------------------------------------------

DLFALLAAFHKSCFQITSPGYALNSMKVWLLEHMKCEIIKEFPEVYFCERPGSFYGTLFT

------------------------------------------------------------

WKQRTPFEGILIIYSRNQTVMFQCLHNLIRILPINCFLKNLKSGSENFLIDNMAFTLEKE

------------------------------------------------------------

LVTLSSLSSAIAKHESNFMQRCEVSKGKSSVVAAALSDRRENIHPYRKELQREKKKMLQT

---------------------------------

NLKVSGALYREITLKVAEVQLKSDFAAQKLSNL
```

**Figure S5. SMURF analysis indicates high probability of a beta-propeller structure in FANCB.** Human FANCB protein sequence was analyzed using SMURF (http://smurf.cs.tufts.edu/). Results shown are for the highest probability alignment of a 6-bladed beta propeller against the SMURFLite library of 207 beta-structural SCOP superfamilies. The human FANCB sequence is shown in capital letters. Vertical lines indicate alignment, BBB etc indicates proposed beta-structure alignment. The design of the SMURF Makkov Random Field leads to a high confidence the motif is present when the *p* value is less than 0.001. For example, 478 of 506 bacterial YYY proteins that contain known beta propeller structures have a p value of <0.01 using SMURF.[11]

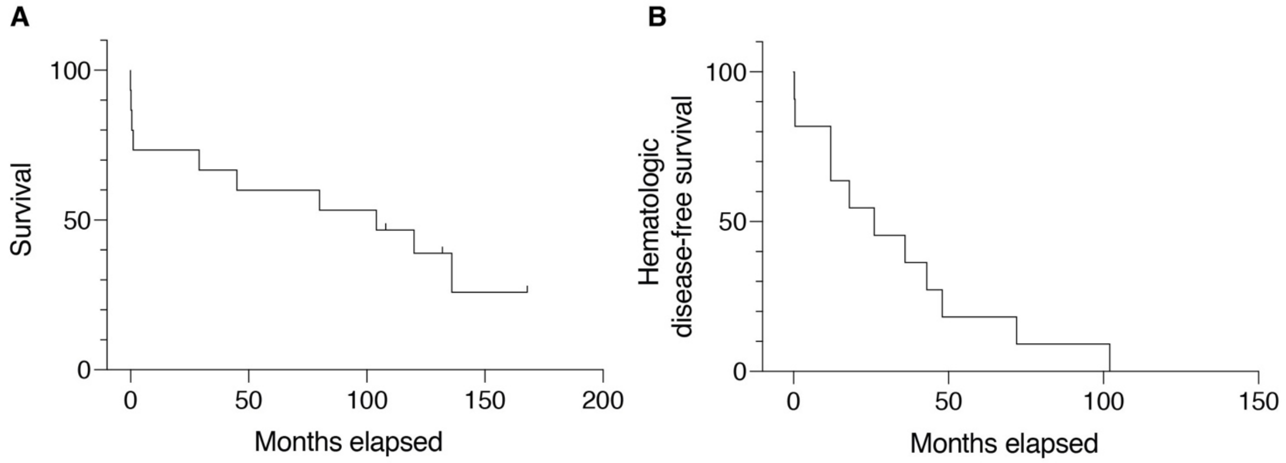

**Figure S6. Time to death and time to hematologic disease for all individuals with *FANCB* variants.**

(A) Kaplan-Meier analysis for time to death including all individuals with *FANCB* variants shows the median time to death of 104 months (n=15; individuals 1, 2, 3, 4, 5, 6, 11, 13, 14, 16, 17, 18, 19, 20 and 21). Individuals excluded from this analysis are detailed in the supplemental methods.

(B) Kaplan-Meier analysis for time to hematologic disease including all individuals with *FANCB* variants shows the median of 26 months (n=11; individuals 3, 5, 6, 11, 14, 16, 17, 18, 19, 20 and 21). Individuals excluded from this analysis are detailed in the supplemental methods.
